# Nuisance Effects and the Limitations of Nuisance Regression in Dynamic Functional Connectivity fMRI

**DOI:** 10.1101/285239

**Authors:** Alican Nalci, Bhaskar D. Rao, Thomas T. Liu

**Affiliations:** Center for Functional MRI, University of California San Diego, 9500 Gilman Drive MC 0677, La Jolla, CA 92093; Department of Electrical and Computer Engineering, University of California San Diego, 9500 Gilman Drive, La Jolla, CA 92093; Departments of Radiology, Psychiatry, and Bioengineering, University of California San Diego, 9500 Gilman Drive, La Jolla, CA 92093

**Keywords:** dynamic functional connectivity, nuisance regression

## Abstract

In resting-state fMRI, dynamic functional connectivity (DFC) measures are used to characterize temporal changes in the brain’s intrinsic functional connectivity. A widely used approach for DFC estimation is the computation of the sliding window correlation between blood oxygenation level dependent (BOLD) signals from different brain regions. Although the source of temporal fluctuations in DFC estimates remains largely unknown, there is growing evidence that they may reflect dynamic shifts between functional brain networks. At the same time, recent findings suggest that DFC estimates might be prone to the influence of nuisance factors such as the physiological modulation of the BOLD signal. Therefore, nuisance regression is used in many DFC studies to regress out the effects of nuisance terms prior to the computation of DFC estimates. In this work we examined the relationship between seed-specific sliding window correlation-based DFC estimates and nuisance factors. We found that DFC estimates were significantly correlated with temporal fluctuations in the magnitude (norm) of various nuisance regressors. Strong correlations between the DFC estimates and nuisance regressor norms were found even when the underlying correlations between the nuisance and fMRI time courses were relatively small. We then show that nuisance regression does not necessarily eliminate the relationship between DFC estimates and nuisance norms, with significant correlations observed between the DFC estimates and nuisance norms even after nuisance regression. We present theoretical bounds on the difference between DFC estimates obtained before and after nuisance regression and relate these bounds to limitations in the efficacy of nuisance regression with regards to DFC estimates.

## 1. Introduction

In resting-state functional magnetic resonance imaging (fMRI), the correlation between the BOLD time courses from different brain regions is used to estimate the functional connectivity (FC) of the brain in the absence of an explicit task. This approach has revealed a number of resting-state networks, where each network consists of brain regions that exhibit a high degree of mutual correlation (Fox et al., 2005). For the most part, FC estimates have assumed temporal stationarity of the underlying BOLD time courses and are obtained by computing correlations over the entire scan duration. As the resulting estimates represent a *temporally averaged* measure of brain connectivity over a typical scan duration of 5–10 minutes, they can miss important dynamic temporal changes in FC (Allen et al., 2014; Preti et al., 2017; Hutchison et al., 2013).

An increasing number of studies have focused on the dynamics of functional brain connectivity by considering dynamic FC (DFC) measures that are computed over time scales typically on the order of tens of seconds and thus much shorter than the scan duration (Allen et al., 2014; Preti et al., 2017; Hutchison et al., 2013). Approaches for estimating DFC include sliding window correlation method (Hutchison et al., 2013; Calhoun et al., 2014), time-frequency methods such as wavelet transform coherence (Chang and Glover, 2010; Yaesoubi et al., 2015), and probabilistic inference methods such as hidden Markov modeling (Vidaurre et al., 2017). To date, the sliding window correlation approach is the most widely used DFC estimation method (Hutchison et al., 2013). In this approach, the correlations between BOLD time courses from different brain regions are computed over sliding windows with durations typically ranging from 30-60 seconds (Preti et al., 2017). Regardless of the analysis method, non-neural processes that affect the BOLD time series can also contaminate the DFC estimates (Murphy et al., 2013; Preti et al., 2017; Hutchison et al., 2013). These confounds are often referred to as *nuisance terms* and include the effects of motion, cardiac and respiratory activity, and fluctuations in arterial CO_2_ concentration. Hutchison et al. (2013) noted that non-neuronal sources that introduce spatial correlations into the time series can also give rise to spurious dynamics in FC measures. Recently, Nikolaou et al. (2016) reported that temporal fluctuations in network degree were related to fluctuations in both heart rate and end-tidal CO_2_. Glomb et al. (2017) found that temporal fluctuations in FC were related to temporal variations in a global measure of average BOLD signal magnitude.

A common step in most fMRI analyses is the use of nuisance regression to minimize the contributions of nuisance terms in BOLD time courses (Liu, 2016; Ciric et al., 2017). Nuisance regressors include cardiac and respiratory activity derived time courses (Birn et al., 2008; Chang et al., 2009), head motion parameters, Legendre polynomials to model scanner drift, signals from white-matter (WM) and cerebrospinal fluid (CSF) regions, and a whole brain global signal (GS) (Liu et al., 2017). Although nuisance regression is widely employed prior to the computation of DFC estimates (Murphy et al., 2013), efforts to examine its efficacy with regards to DFC estimates have been limited. Hutchison et al. (2013) noted that residual nuisance effects “inevitably remain” in the BOLD time series and must be considered in the interpretation of DFC measures. In support of this view, Nikolaou et al. (2016) found that nuisance regression diminished but did not completely remove the relationship between network degree and measures of cardiac and respiratory activity. On the other hand, Xu et al. (2018) reported that global signal regression (GSR) had a spatially heterogeneous impact on DFC estimates, but did not assess whether GSR removed GS contributions from the DFC estimates.

In this paper, we take a closer look at (1) the role of nuisance terms in correlation-based DFC estimates and (2) the efficacy of nuisance regression for DFC studies. In particular, we use two independent datasets to examine the relationship between seed-specific sliding window correlation-based DFC estimates and the norms of various nuisance regressors (WM and CSF signals, GS, cardiac and respiratory measurements, and head motion). We then assess the effect of nuisance regression on this relation, considering both regression applied to the entire scan and regression applied on a sliding window basis. To interpret the empirical findings, we derive mathematical expressions to describe the effect of regression on DFC estimates and compare the experimental results with the theoretical predictions. Preliminary versions of this work have been presented in (Nalci et al., 2017a; Nalci and Liu, 2018).

## 2. Methods

### 2.1. Datasets

In this work, we analyzed two datasets in order to show the generality of our results and to experiment with different types of nuisance measurements. First, to understand the effect of nuisance measurements derived directly from the MRI images such as the GS, WM and CSF, and head motion (HM) time courses on the DFC estimates, we used a publicly available dataset originally analyzed by Fox et al. (2007), which we will refer to as the BS002 dataset. Second, to understand the effect of physiological measurements such as changes in the respiration and cardiac rate on the DFC estimates, we used the dataset analyzed by Wong et al. (2012), which we will refer to as the CFMRI dataset.

BS002 data were acquired from 17 young adults (9 females) using a 3 T Siemens Allegra MR scanner. Each subject underwent 4 BOLD-EPI fixation runs (32 slices, TR=2.16 s, TE=25 ms, 4×4×4 mm), each lasting 7 minutes (194 frames). The subjects were instructed to look at a cross-hair and asked to remain still and awake. High-resolution T1-weighted anatomical images were acquired for the purpose of anatomical registration (TR=2.1 s, TE=3.93 ms, flip angle=7 deg, 1×1×1.25 mm).

CFMRI data were acquired from 10 healthy volunteers (4 males and 6 females) using a 3 Tesla GE Discovery MR750 system. Each subject underwent four separate resting-state scans (5 minutes per scan with either their eyes open or closed). High resolution anatomical data were collected using a magnetization prepared 3D fast spoiled gradient (FSPGR) sequence (TI=600 ms, TE=3.1 ms, flip angle = 8 degrees, slice thickness = 1 mm, FOV = 25.6 cm, matrix size = 256×256× 176). Whole brain BOLD data were acquired over thirty axial slices using an echo planar imaging (EPI) sequence (flip angle = 70 degrees, slice thickness = 4 mm, slice gap = 1 mm, FOV = 24 cm, TE = 30 ms, TR = 1.8 s, matrix size = 64×64). Cardiac pulse and respiratory data were monitored using a pulse oximeter (InVivo) and a respiratory effort transducer (BIOPAC), respectively. The pulse oximeter was placed on each subject’s index finger while the respiratory effort belt was placed around the subject’s abdomen. Physiological data were sampled at 40 Hz using a multi-channel data acquisition board (National Instruments).

### 2.2. Preprocessing steps for the BS002 dataset

Standard pre-processing steps were conducted with the AFNI software package (Cox, 1996). The initial 9 frames from each EPI run were discarded to minimize longitudinal relaxation effects. Images were then slice-time corrected and co-registered, and the 6 head motion parameter time series were retained. The resultant images were converted to coordinates of Talairach and Tournoux (TT), resampled to 3 mm cubic voxels, and spatially smoothed using a 6 mm full-width-at-half-maximum isotropic Gaussian kernel. The 1^*st*^ and 2^*nd*^ order Legendre polynomials (a constant term to model the temporal mean and a linear trend) were projected out from each voxel’s time course. Each voxel time series was then converted into a percent change BOLD time series through demeaning and division by its mean value.

To analyze the effect of nuisance regression on DFC estimates, we used both (1) full linear regression in which regression is performed over the whole scan duration and (2) block regression (also referred to as sliding window regression) in which regression is performed for each window separately (more details on block regression are provided in Section 2.6). For each type of regression, we performed a separate regression on the data using one of the following types of regressor: (1) the set of all 6 head motion parameters, (2) nuisance signals from WM and CSF regions, and (3) the global signal (GS), which was calculated as the average of the percent change time series across all voxels within the brain.

For DFC analysis, we used seed signals derived from the posterior cingulate cortex (PCC), intraparietal sulcus (IPS), frontal eye fields (FEF), and motor network (MOT). These seed signals were obtained by averaging time series selected over spheres of radius 6 mm (2 voxels) centered about their corresponding TT coordinates (He and Liu, 2012). The sphere centers were obtained by converting the MNI coordinates from Van Dijk et al. (2010) to TT coordinates (Lacadie et al., 2008). For the PCC, left MOT, and right MOT seeds we used the coordinates [0,−51,26], [−36,−22,52] and [37,−21,52], respectively. A combined MOT seed was obtained by using the left and right MOT coordinates to define two spheres and by merging them. For the IPS and FEF seeds, we used the coordinates [27,-58,49] and [24,−13,51] from (Fox et al., 2006). Finally, for the WM and CSF nuisance signals, we defined the sphere centers as [12,−36,27] and [9,−9,15], respectively.

### 2.3. Preprocessing steps for the CFMRI dataset

We used FSL and AFNI packages to preprocess the CFMRI dataset (Woolrich et al., 2009; Smith et al., 2004). First, high-resolution anatomical data were skull stripped and segmentation was applied to estimate WM, gray matter, and CSF partial volume fractions. Images were then slice-time corrected and co-registered and the 6 head motion parameter time series were retained. The anatomical data were then aligned to the functionals. Each subject’s native space data were transferred to TT coordinates and spatially smoothed using a 6 mm full-width-at-half-maximum isotropic Gaussian kernel. A binary brain mask was created for each subject using the transferred data and was eroded by 2 voxels along the brain edges to eliminate possible inclusion of non-brain areas (Rack-Gomer and Liu, 2012). The first 6 frames (10.8 seconds) from each functional data were discarded to minimize longitudinal relaxation effects. For each run, the 1^*st*^ and 2^*nd*^ order Legendre polynomials and the 6 head motion parameters were projected out to obtain the ‘baseline’ CFMRI data.

We performed RVHRCOR by simultaneously projecting out the physiological nuisance regressors derived from cardiac and respiratory signals using both full linear regression and block regression (Chang et al., 2009; Birn et al., 2008). Specifically, a respiratory variation (RV) signal was computed as the standard deviation of the respiratory signal using a 7.2 second sliding window and was then convolved with the respiration response function (RRF) to obtain the respiration regressor (RVf). Similarly, a heart rate (HR) signal was computed as the inverse of average peak to peak time interval between two consecutive heartbeats over a 7.2 second sliding window. The HR was then convolved with a cardiac response function (CRF) to obtain the cardiac regressor (HRf). In this process, we discarded 4 scans out of 40 due to the poor quality of physiological signals, leaving a total of 36 scans for analysis. For DFC analysis, we used the same seed coordinates as mentioned for the BS002 dataset.

### 2.4. Calculation of the DFC estimates: Sliding window correlations

The DFC estimates were obtained by computing the sliding window correlations between pairs of seed signals selected from the PCC, IPS, FEF, and MOT regions. Specifically, we computed the sliding window correlations between the following seed pairs: PCC and IPS, PCC and FEF, PCC and MOT, IPS and FEF, IPS and MOT, and FEF and MOT. Denoting *x*_1_ and *x*_2_ as two seed time series and *x*_1,*k*_ and *x*_2,*k*_ as the windows taken from those signals (where *k* denotes the window index), we computed the window correlation value as 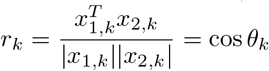. Here, *θ_k_* is the observed angle between those time courses and |·| is the vector norm. Note also that here and throughout the paper the time courses are represented as column vectors. This process was repeated by temporally shifting the window index until all sliding window correlations were carried out over the entire scan duration. The final aggregated set of correlation values *r_DFC_* = [*r*_1_,*r*_2_, · · ·, *r_T_*] was used as an estimate for the DFC.

In our primary analysis, to maximize the number of available DFC samples per scan, the sliding window duration in this paper was fixed at 30 TRs and a window shift of 1 TR was used (Hutchison et al., 2013; Xu et al., 2018). This duration corresponds to a window length of 63 seconds for the BS002 dataset and 54 seconds for the CFMRI dataset. These window lengths lie within the range of durations typically used in the existing DFC literature (Preti et al., 2017). In supplementary analyses, we considered two additional window sizes: a longer window of duration of 100 seconds, which corresponds to 48 TRs for the BS002 dataset and 56 TRs for the CFMRI dataset, and a shorter window size of 40 seconds, which corresponds to 20 TRs for the BS002 dataset and 22 TRs for the CFMRI dataset.

### 2.5. Calculation of the nuisance metrics: Nuisance norms

To examine the relationship between the nuisance measurements and the DFC estimates, we used the norm of the regressor as our nuisance metric. Note that the norm of a regressor is simply the geometric length of the column vector used to represent the regressor time course. In Section 4 we present some toy examples and geometric arguments to show how DFC estimates can be related to the norms of various nuisance terms. The theoretically inclined reader may choose to read that section first before continuing.

For a single regressor, the computation of the windowed norms was done by first demeaning the nuisance regressor for each window. Denoting *n_k_* as the demeaned nuisance measurement in the *k*th window, the window norm was computed as 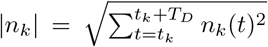, where *t* indicates time in TR units and *T_D_* is the window duration minus 1 (e.g. *T_D_* = 29 for a window duration of 30 TRs). When multiple regressors were used in the regression (e.g. for HM), we computed the total norm as 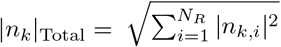, where *i* denotes the index for different regressors and *N_R_* is the number of regressors. In the text we use the term norm and the notation |*n_k_*| to denote both |*n_k_*| and |*n_k_*|_Total_. The norm was computed for all windows by temporally shifting the window index. The final set {|*n*_1_|, |*n*_2_|,…, |*n_T_*|} of nuisance norms comprised the nuisance norm time course. To examine the relationship between the nuisance norm and DFC estimates, we computed the correlation coefficient between the DFC estimates and the nuisance norm time courses.

### 2.6. Analysis of nuisance regression techniques

We applied two types of linear regression, which we refer to as full regression and block regression. In full regression, the nuisance terms were projected out of the voxel time series over the entire scan duration. For example, for a single regressor, the clean time course after full regression was obtained as *x̃* = *x* − *n*(*n^T^n*)^−1^*n^T^x* = *x* − *bn*, where *n* is the column vector representing the nuisance regressor and *b* = (*n^T^n*)^−1^*n^T^x* is the scalar fit coefficient for the entire scan duration. In block regression, the nuisance measurement was projected out of each window separately, such that the clean time course for the *k*th window was 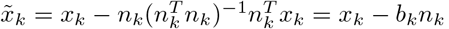, where 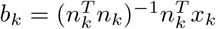 is the window-specific scalar fit coefficient.

Note that we will refer to the “clean” DFC estimates obtained after the application of regression as Post FullReg DFC and Post BlockReg DFC estimates for full and block regression, respectively. To simplify the presentation, we will also use the shorter term Post DFC to refer to Post FullReg DFC, since full regression is the method that has typically been used in the literature.

### 2.7. Significance testing of the relationship between DFC estimates and nuisance norms

To assess the statistical significance of the relationship between the DFC estimates and nuisance norms, we used an autoregressive (AR) bootstrapping procedure based on the work of (Chang and Glover, 2010; Efron and Tibshirani, 1986; Zalesky et al., 2014) As our null hypothesis was that the DFC estimates and nuisance norms were not linearly related, we fit (on a per scan basis) separate AR processes of model order *q* to the DFC estimates and nuisance norm time courses both before and after full/block regression. The model order *q* was determined according to the Bayesian information criterion (BIC). Using the estimated AR coefficients from each scan, we generated 10,000 surrogate time series for both the DFC estimates and nuisance norms and then computed the associated surrogate correlation coefficients.

To create a null distribution for the assessment of significance across the study sample, we computed the absolute value of each of the surrogate correlation coefficients in the sample and took the mean of the absolute values across the sample. The resulting null distribution consisted of 10,000 surrogate mean absolute correlation values. We used this null distribution to assess the significance of the sample mean absolute correlation value.

As a secondary descriptive analysis, we also assessed significance on a per-scan basis. We formed a null distribution of correlation values for each scan by correlating the surrogate nuisance norms with the surrogate DFC estimates. We used the resulting null distribution to compute the p-value associated with the correlation coefficient computed from each scan’s measured data. We then calculated the percentage of scans that showed significant correlations between the DFC estimates and nuisance norms for significance thresholds of 0.05 and 0.10.

## 3. Results

In this section, we first show that the DFC estimates obtained between pairs of seed time courses can be significantly correlated with the norms of various nuisance measurements. We demonstrate that strong correlations between the DFC estimates and nuisance norms exist even when the correlations between the nuisance and seed time courses are small. We then show that performing nuisance regression prior to the computation of the DFC estimates does not necessarily eliminate the presence of strong and significant correlations between the nuisance norms and DFC estimates.

**Figure 1:**
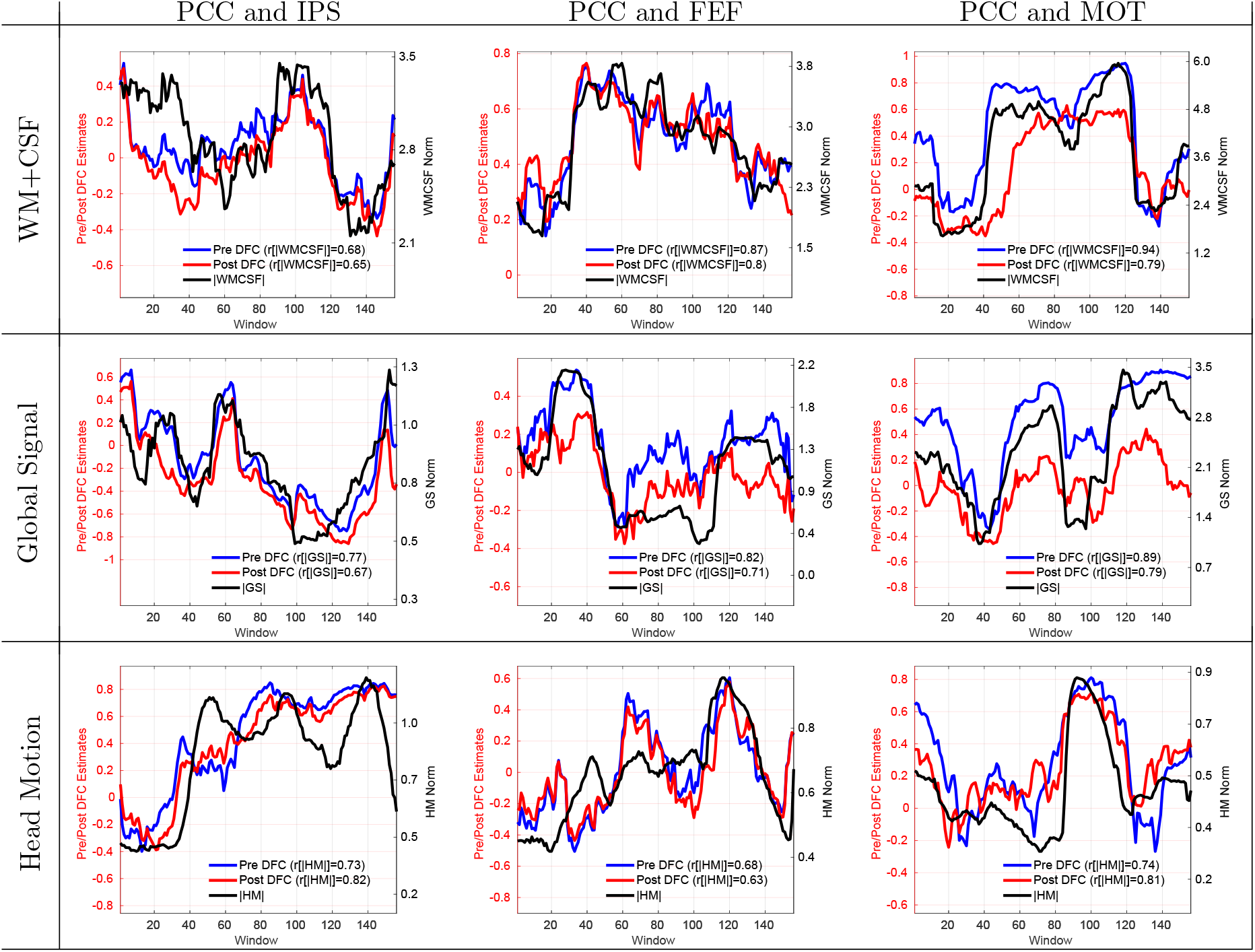
9 example scans that demonstrate strong and significant (*p* < 0.02) correlations between the nuisance norms and DFC estimates. The type of nuisance regressor is indicated by the row label. The seed pair for the DFC estimate is indicated by the column label. The solid blue line in each panel shows the DFC estimate prior to nuisance regression (labeled as Pre DFC on the legend) and the solid black line shows the corresponding nuisance norm. Correlation values between the DFC estimates and nuisance norms are indicated in the legend labels. The correlations between the nuisance norms and the Pre DFC estimates varied from *r* = 0.68 to *r* = 0.94 with a mean value of *r* = 0.79. The DFC estimates after performing full linear regression are shown with solid red lines (labeled as Post DFC on the legend). The correlations between the nuisance norms and the Post DFC estimates ranged from *r* = 0.63 to *r* = 0.82 with a mean value of *r* = 0.74.

### 3.1. Examples of correlations between DFC estimates and nuisance norms

In Figure 1, we present examples of scans from the BS002 dataset in which significant positive correlations between the DFC estimates and nuisance norms were observed. The column labels indicate the seed region pair (e.g. PCC and IPS, PCC and FEF, and PCC and MOT) and the row labels indicate the type of nuisance norm. The solid blue line in each panel shows the DFC estimate before nuisance regression (labeled as Pre DFC) and the solid black line shows the respective nuisance norm. In these scans, the correlations between the nuisance norms and the Pre DFC estimates ranged from *r* = 0.68 to *r* = 0.94 with a mean correlation value of *r* = 0.79, corresponding to a range of explained variances from 46% to 88% over the example scans. The nuisance norms in these scans were significantly correlated with the DFC estimates with per-scan p-values across the example scans of *p* < 0.0075 for WM+CSF, *p* < 0.0009 for GS and *p* < 0.0237 for HM regressors.

**Figure 2:**
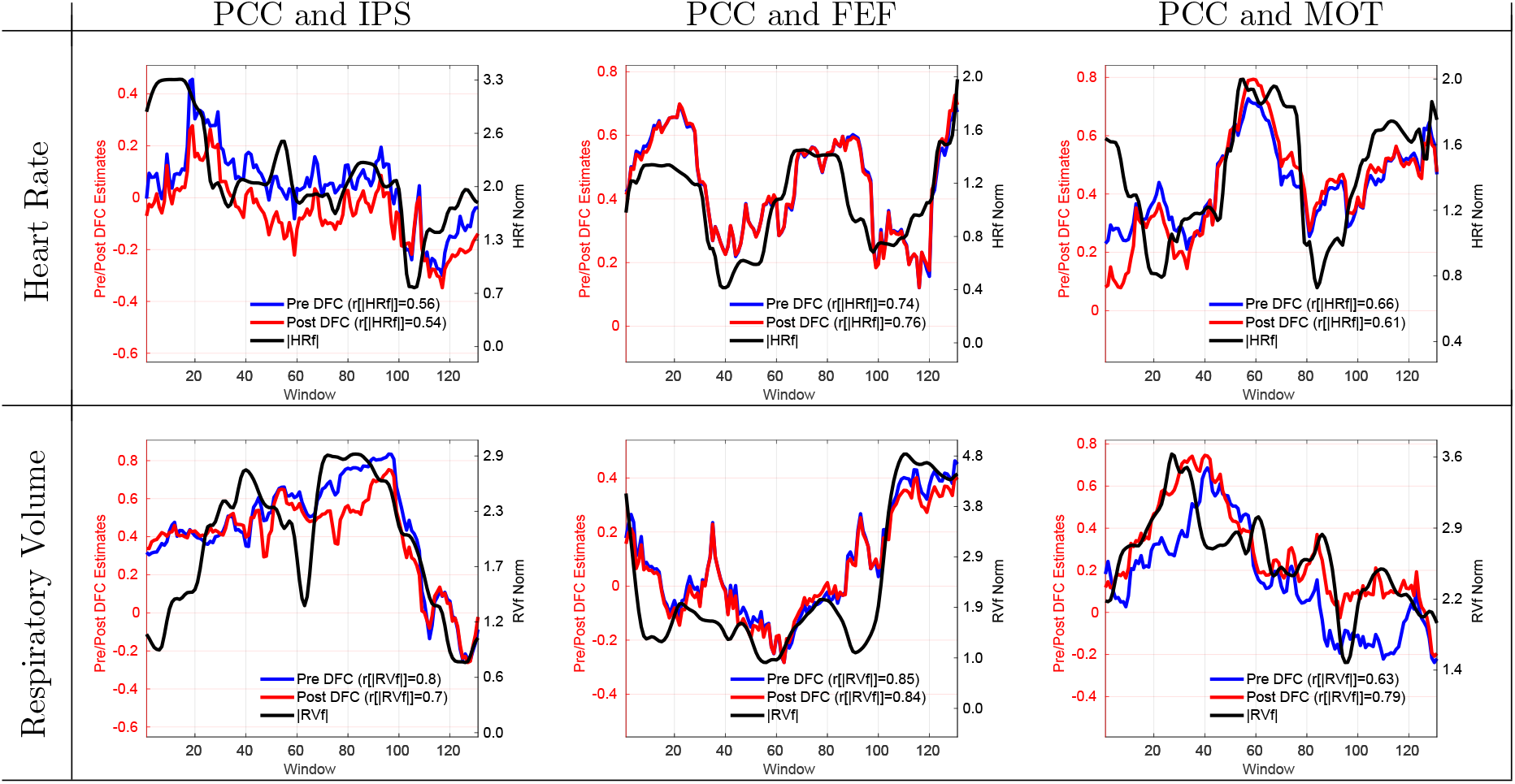
6 example scans showing that the DFC estimates are significantly (*p* < 0.03) correlated with the norms of physiological nuisance terms. Row labels indicate the type of physiological nuisance term considered and column labels indicate the seed pair. Correlation values between the DFC estimates and nuisance norms are indicated in the legend labels. The correlations between the Pre DFC estimates and the nuisance norms ranged from *r* = 0.56 to *r* = 0.85 with a mean correlation of *r* = 0.71. The DFC estimates obtained after performing RVHRCOR (Post DFC) are similar to the Pre DFC estimates. The correlations between the Post DFC and the nuisance norms ranged from *r* = 0.54 to *r* = 0.84 with a mean correlation of *r* = 0.71.

In Figure 2, we show 6 example scans from the CFMRI dataset demonstrating significant positive correlations between DFC estimates and physiological nuisance norms. The first row shows 3 scans using the HRf norm as the nuisance measure and the second row shows 3 scans using the RVf norm. Each column shows a different seed pair for the DFC estimates. The correlations between the Pre DFC estimates and the nuisance norms in these scans ranged from *r* = 0.56 to *r* = 0.85 with a mean correlation of *r* = 0.71 corresponding to a range of explained variances from 31% to 72%. The RVf and HRf norms were significantly correlated with the DFC estimates with per-scan p-values of *p* < 0.0345 for RVf and *p* < 0.0355 for the HRf regressors.

**Figure 3:**
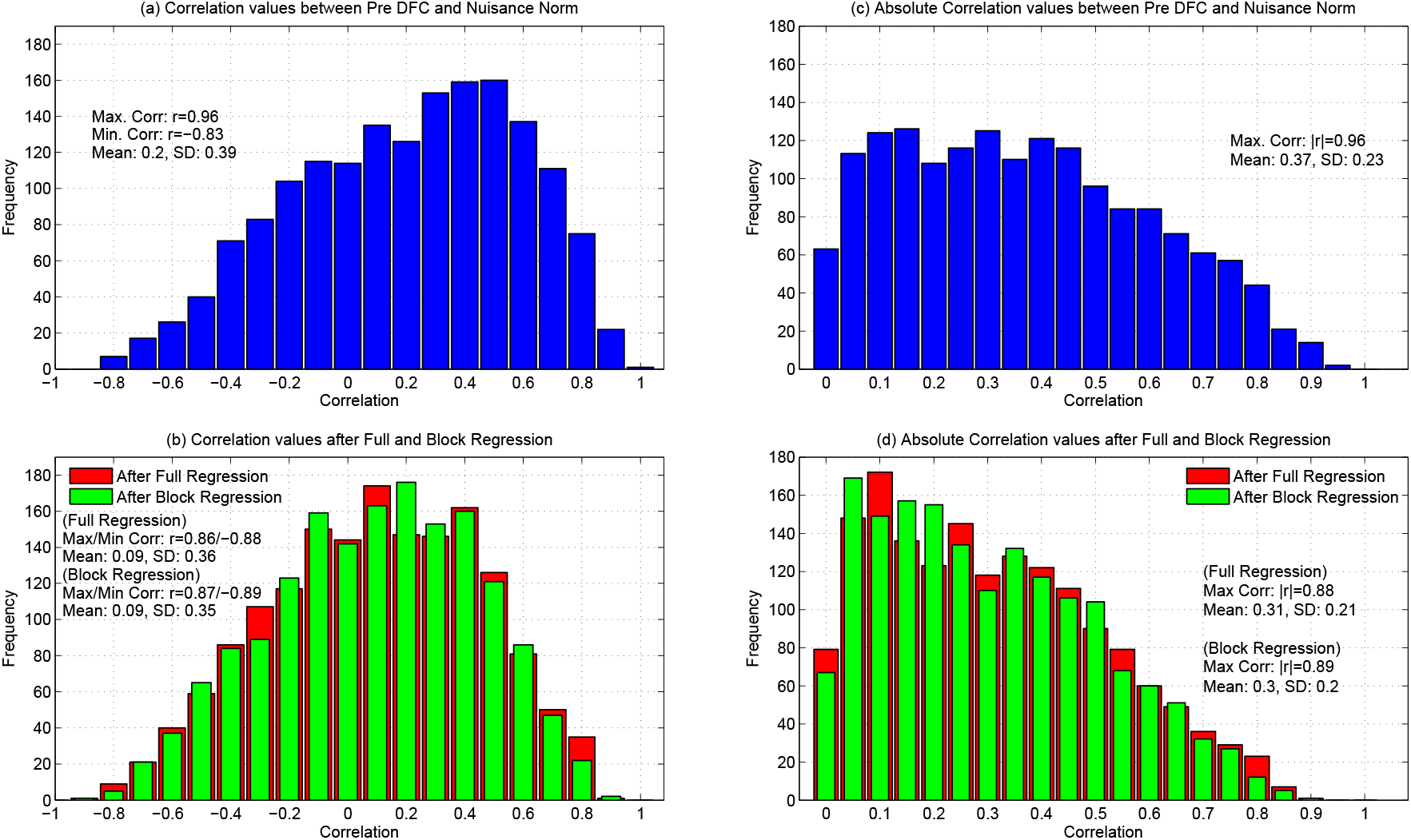
(a) Histogram of correlation values obtained between the Pre DFC estimates and the nuisance norm across all scans, seed pairs, and nuisance norms. Distribution of the correlation values ranged from *r* = −0.83 to *r* = 0.96. (b) After regression, the correlation values ranged from a minimum of *r* = −0.88 to a maximum of *r* = 0.86 for full regression, and from a minimum of *r* = −0.89 to a maximum of *r* = 0.87 for block regression. (c) Histogram of absolute correlation values between the nuisance norm and Pre DFC estimates with a sample mean of |*r*| = 0.37. (d) Absolute correlation values after full and block regression with sample means of |*r*| = 0.31 for full and |*r*| = 0.30 for block regression.

In Supplementary Figures 1 and 2 we show additional examples for each regressor and seed pair in cases of medium to weak positive correlations observed between the DFC estimates and nuisance norms. Note that all qualitative examples shown in this paper are from different scans but can be from the same subject.

In Figure 3a, we show the histogram of correlations between the nuisance norms and the DFC estimates across all scans, seed pairs, and nuisance norms. The histogram includes a total of 1, 656 correlation values with 1, 224 belonging to the BS002 dataset (68 scans × 6 seed pairs × 3 types of nuisance norm) and the remaining 432 from the CFMRI dataset (36 scans × 6 seed pairs × 2 types of nuisance norm). The correlations ranged from a negative value of *r* = −0.83 to a positive value of *r* = 0.96, with a skewed distribution in which 68% of the correlations were positive and the remaining 32% were negative. In Supplementary Figure 3, we present the correlations from Figure 3a separated into the values for individual regressors. Figure 3c shows the histogram of absolute correlation values, with a sample mean value of |*r*| = 0.37. The significance of the sample mean absolute correlation value is assessed in the following section.

In Figure 4, we show 3 example scans to demonstrate the existence of significant anti-correlations between the Pre DFC estimates and various regressor norms. The type of nuisance regressor and the seed region pair are indicated in the column labels. For these scans, the correlation values were strongly negative ranging from *r* = −0.82 to *r* = −0.61 with an increase in nuisance norm corresponding to a decrease in the DFC estimate. The nuisance norms in these scans were significantly anti-correlated with the DFC estimates with per-scan p-values of *p* < 0.001 for HRf, *p* < 0.0024 for WM+CSF and p < 0.0226 for the HM regressors.

**Figure 4:**
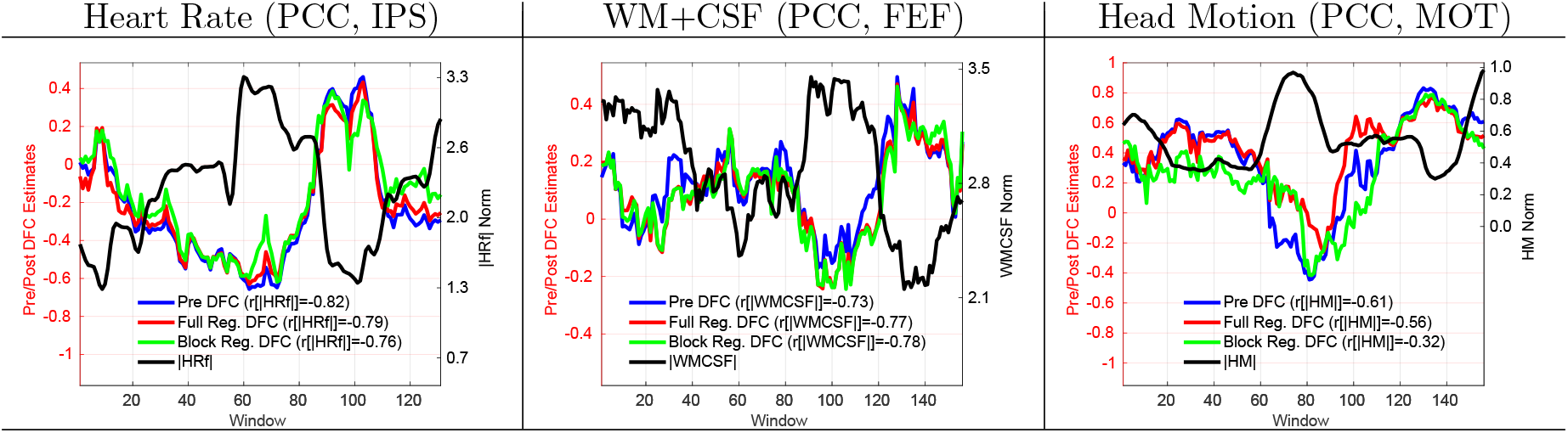
3 scans showing examples of strong anti-correlations between the nuisance norms and the DFC estimates both before and after regression. The type of nuisance regressor and the seed region pair are indicated in the column labels. The correlation values between the DFC estimates and the nuisance norms are indicated in the legend labels.

As an exploratory analysis, we also assessed whether the state of the eyes (open or closed) in the CFMRI dataset affected the relationship between the DFC estimates and the norms of the RVf and HRf time courses. We found no significant difference (*p* = 0.95, two-tailed paired t-test) in the correlations obtained between eyes open and eyes closed conditions.

### 3.2. Assessing significance across the sample

We now consider whether the DFC estimates are significantly correlated with nuisance norms across the study sample. As demonstrated in the previous section, both positive and negative correlations can exist between the DFC estimates and nuisance norms. Taking this into account, we focused on the absolute value of the correlation values and computed the mean of the absolute correlation values across the sample. As described in the Methods section, we assessed the significance of the mean absolute value using an empirical null distribution. Figure 5a shows the empirical null distribution and the sample mean absolute value, which was found to be significant with *p* < 10^−6^. Note that the sample mean absolute value indicated here is simply the mean of the histogram of the absolute correlation values shown in Figure 3c. In panels (b) and (c), we show the null distributions and sample values when considering complementary subsets of the sample where the global signal (GS) is either included or not included, respectively, with associated p-values of *p* < 10^−6^ for both subsets. In Supplementary Figure 4, similar plots are provided for analyses with window lengths of approximately 40 and 100 seconds.

**Figure 5:**
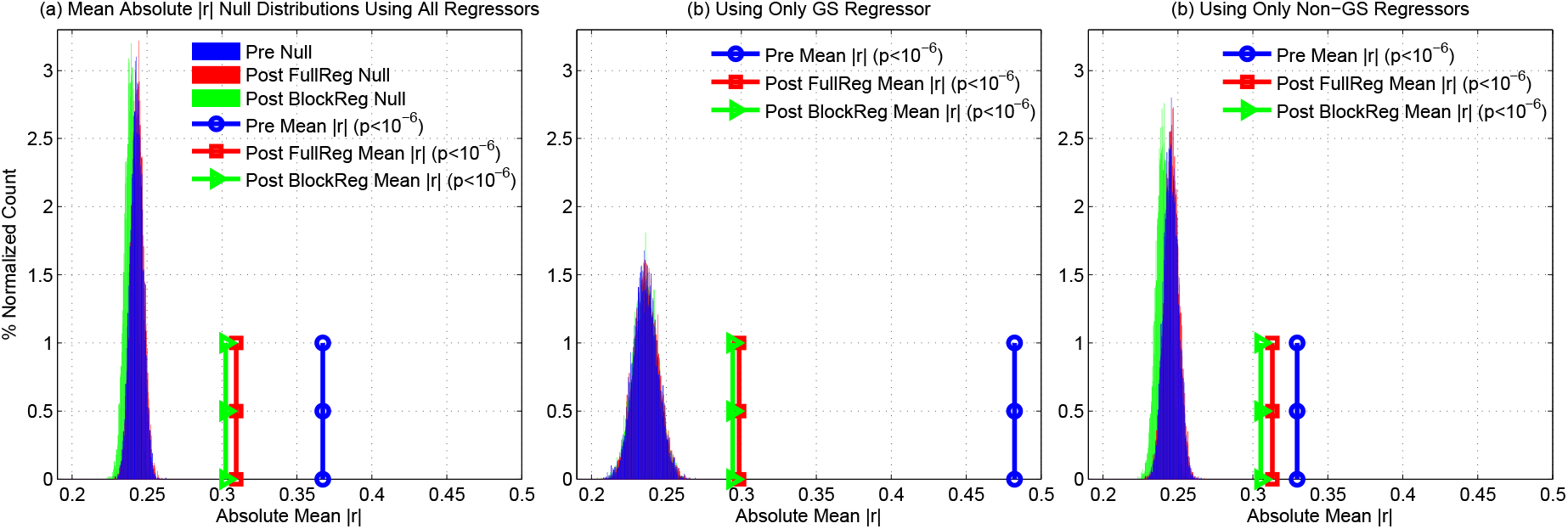
(a) Empirical null distributions for mean absolute correlation values when considering all regressors both before regression (blue) and after full regression (red) or block regression (green). The sample mean absolute correlation values for these conditions are shown by the circles, squares, and triangles, and are equal to the values previously reported in 3c-d. The sample mean absolute correlation values were found to be significant (*p* < 10^−6^) both before and after full or block regression. (b) Empirical null distributions and sample mean absolute correlation values when considering only the GS regressor. There was a marked reduction in the sample mean absolute correlation values after regression. However, the sample mean absolute correlation values were still found to be significant (*p* < 10^−6^) both before and after nuisance regression. (c) Empirical null distributions and sample mean absolute correlation values for the Non-GS regressors. Sample mean absolute correlation values were found to be significant (*p* < 10^−6^) both before and after nuisance regression.

To complement our primary analysis, we also considered the extent to which the correlation coefficients were significant at the per-scan level. Using the scan-specific null distributions (see Methods), we found that 24% of the correlations between the nuisance norms and Pre DFC estimates were significant at the *p* < 0.05 level and 30% of the correlation values were significant at the *p* < 0.10 level. In supplementary material, Table 1 summarizes these findings and provides results for analysis window lengths of 40s and 100s. The results for a shorter window length of 40s were fairly similar to that of our primary results (approximate window length of 60s). However, we observed a slightly higher percentage of significance scans at the longer window length of 100s.

### 3.3. Strong correlations between DFC estimates and nuisance norms exist even when the underlying correlations between the nuisance and seed time courses are small

In this section, we ask the question: can strong correlations between DFC estimates and nuisance norms occur even when the nuisance time course are only weakly correlated with the seed time courses? Figure 6a shows the correlations between the DFC estimates and the nuisance norms plotted against the root-mean-squared (RMS) correlations between the underlying nuisance and seed time courses. Each point corresponds to a specific scan, nuisance term, and seed region pair. For each point, the RMS correlation is computed by correlating the nuisance time course with each of the two seed time courses (from the same scan) and then calculating the RMS value (e.g. 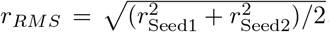, where *r*_Seed1_ and *r*_Seed2_ are the correlation coefficients between the nuisance time course and the first and second seed signals, respectively).

We first observe that strong correlations (points above and below the dashed lines in Figure 6a) between the DFC estimates and nuisance norms exist even when the RMS correlation between the nuisance and seed time courses is close to zero. We defined strong correlations between the DFC estimates and nuisance norm as those correlations for which |*r*| > 0.5, corresponding to an explained variance of greater than 25%. This threshold corresponded to an empirical *p* = 0.09 as assessed with the null distribution obtained by correlating the surrogate time series across all scans and realizations. Note that the threshold we have used merely serves as a convenient way to delineate strong correlations and its exact value is not critical. Conclusions similar to those stated here and below would be obtained even if we increased or decreased the threshold.

For further analysis, we binned the points according to the RMS correlation values using a bin width of 0.05. For each RMS correlation bin, we counted the number of points in which the nuisance norm explained more than 25% variance in the DFC estimates (i.e. number of scans above the threshold) and then divided this count by the total number of points across all bins. In Figure 6b, the blue bars indicate the percentage of strong correlations between the DFC estimates and nuisance norms as a function of the binned RMS correlation values. The green bars indicate the percentage of strong correlations associated with the GS regressor. The cumulative percentage of strong correlations versus RMS correlation value is shown by the red dotted line.

**Figure 6:**
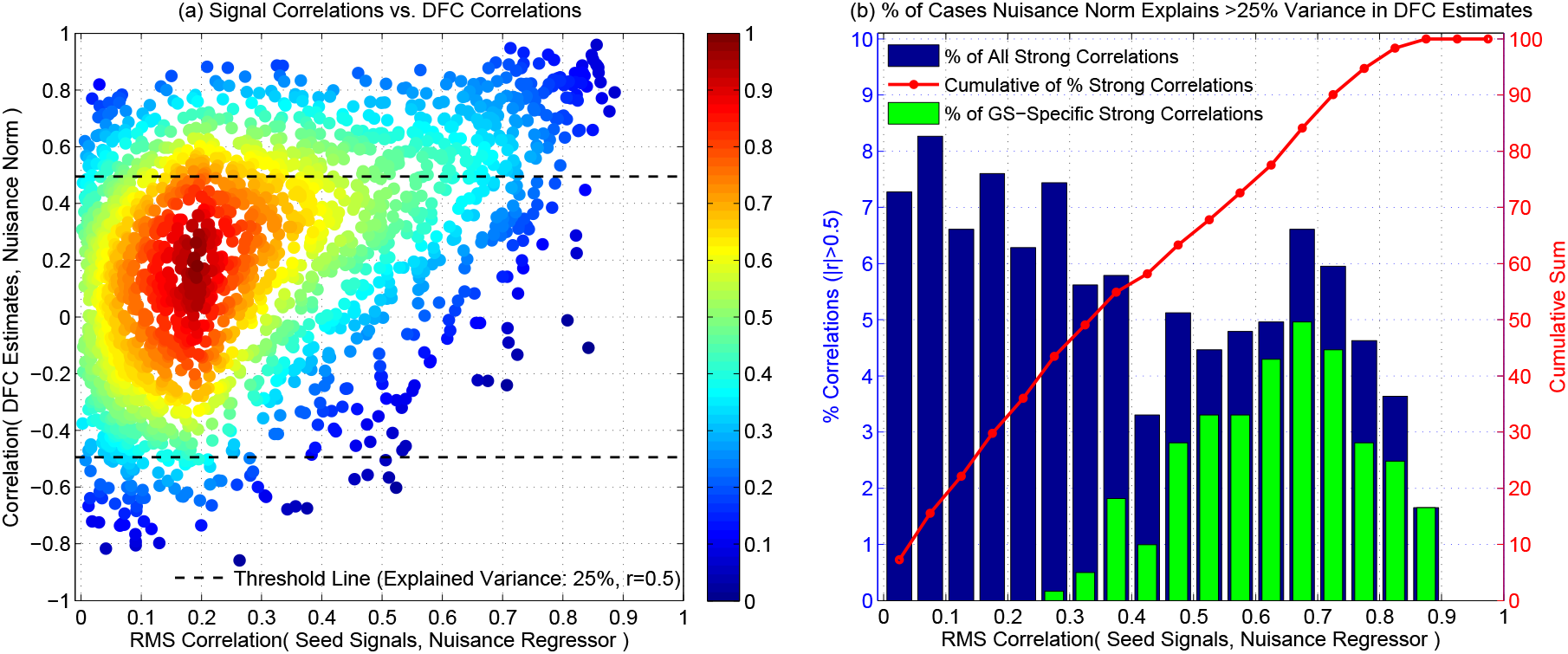
(a) Correlation between the DFC estimate and nuisance norm versus the RMS correlation between the raw nuisance signal and the two seed time courses. Correlation thresholds are indicated by the black dashed lines such that points that lie above and below these lines correspond to cases in which the DFC estimates are considered to be ‘strongly correlated’ (|*r*| > 0.5 with empirical *p* ≤ 0.09) with the nuisance norms such that nuisance norms explain more than 25% of the total variance in the DFC estimates. The relative density of correlations (maximum density is normalized to 1.0) is indicated by the color map on the right-hand side. The relative density is computed by summing the total number of data points in smaller sub-grids and normalizing by the total number of points. (b) The blue bars show the percentage (%) of cases in which the nuisance norms explain more than 25% of the total variance in the DFC estimates as a function of the RMS correlation between the raw nuisance and the two seed time courses. The red dotted line (dots are located at bin centers) shows the cumulative sum of the percentage of strong correlations as a function of the RMS correlations. The green bars show the percentage (%) of strong correlations that are accounted for by the GS regressor.

We found that a large percentage of the strong correlations between the DFC estimates and nuisance norms occurred for fairly small RMS correlation values between the raw nuisance time course and seed signals. For example, 22% of the all strong correlations between the DFC estimates and nuisance norms occurred for RMS values in the interval *r* = 0 to *r* = 0.15. Note that in this interval, the nuisance terms account for less than 2.2% of the average variance of the seed time courses. Excluding the GS, 33% of the strong correlations between the DFC estimates and Non-GS nuisance norms occurred in the same interval. If we expand the interval *r* = 0 to *r* = 0.25, then 36% of the strong correlations lie within this expanded interval, corresponding to nuisance terms that account for less than 6.25% of the average variance of the seed time courses. Excluding the GS, 54% of the strong correlations lie within the same interval. In other words, for 36% of the all cases and 54% of the Non-GS cases examined, the nuisance norm explains more than 25% of the variance in the DFC estimates even though the raw nuisance time courses explain less than 6.25% of the average variance of the seed time courses.

Overall, we have the rather surprising observation that strong correlations between the DFC estimates and nuisance norms can exist even when the underlying nuisance terms account for only a small fraction of the variance of the seed time courses. We present a plausible explanation for this observation in Section 4.3 and examine its implications for nuisance regression later in the text.

Finally, the green bars in Figure 6b show the percentage of strong correlations between the DFC estimates and nuisance norm due to the GS regressor. This shows that the RMS correlation values between the GS and seed signals are relatively higher as compared to the Non-GS regressors. This is expected since the GS is computed as the average of all voxels time courses and is expected to correlate well with the seed time courses. On the other hand, the Non-GS regressors (difference between the blue and green bars) have fairly small RMS correlation values with the seed time courses even though their norms can be strongly correlated with the DFC estimates.

### 3.4. Regression does not eliminate the relation between the DFC estimates and nuisance norms

In this section, we show that full regression does not necessarily eliminate the relationship between the DFC estimates and the nuisance norms. Revisiting Figures 1 and 2, our focus now is on the solid red lines (Post DFC), which show the DFC estimates obtained after projecting out the relevant nuisance regressors. The fluctuations in the Pre DFC and Post DFC estimates in both figures are similar to each other and there is very little difference in their relationships to the nuisance norm. In the scans shown in both figures, the mean correlation value between the Post DFC estimates and the nuisance norms was *r* = 0.73, which is very close to the mean correlation value of *r* = 0.76 between the Pre DFC estimates and the nuisance norms. For these example scans, the Post DFC estimates were still significantly correlated with the nuisance norms after nuisance regression for the WM+CSF, GS and HM regressors (*p* < 0.02) and also for the RVf and HRf regressors (*p* < 0.04).

**Figure 7:**
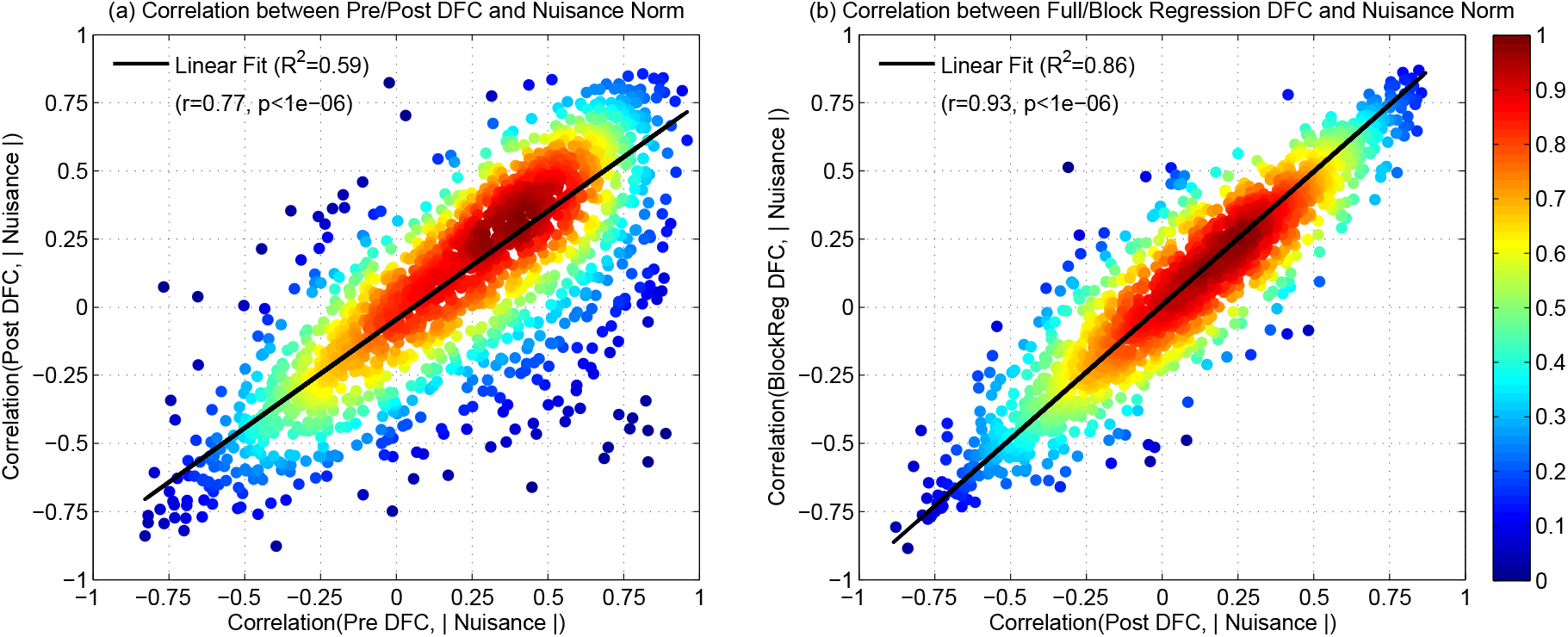
(a) The correlations between the Post DFC estimates and the nuisance norms (y-axis) versus the correlations between the Pre DFC estimates and the nuisance norms (x-axis). Post DFC estimates were still largely correlated with the nuisance norms and there was a strong linear relationship when compared to the correlations between the Pre DFC and nuisance norms (linear fit shown with black solid line, *R*^2^ = 0.59, *r* = 0.77, *p* < 10^−6^). The relative density of correlations (normalized to 1.0) is indicated by the color map on the right hand side. (b) The correlations between the DFC estimates and the nuisance norms after block regression (y-axis) and after full regression (x-axis). There was a strong linear relationship between the correlation populations (fit shown with black solid line, *R*^2^ = 0.86, *r* = 0.93, *p* < 10^−6^). The correlation distributions for full and block regression were not significantly different (paired two-tailed t-test *p* = 0.86). The effect size (*d* = 0.0017) and the absolute difference in correlation population means (0.0006) were negligibly small.

Viewed across the entire sample, the Post DFC correlation values shown in Figure 3b ranged from a minimum of *r* = −0.88 to a maximum *r* = 0.86 with a skewed distribution in which 60% of the correlations were positive and the remaining 40% were negative. The correlation distributions were similar to the Pre DFC histograms in Figure 3a,b with a cosine similarity values of *S* = 0.77 and *S* = 0.87 for the correlation and absolute correlation values, respectively.

Over the sample, the mean absolute correlation |*r*| was found to be significant with *p* < 10^−6^ (see Figure 5a). Moreover, we found that 14% of the correlations between the nuisance norms and Post DFC estimates were still significant at the *p* < 0.05 level and 22% of the correlation values were significant at the *p* < 0. 10 level.

To further demonstrate the relationship between the Pre and Post DFC estimates, in Figure 7a we plotted the correlations between the Post DFC estimates and nuisance norms versus the corresponding correlations between the Pre DFC estimates and the nuisance norms across all scans, nuisance time courses, and seed pairs. The relative density of correlation values is indicated with the color bar on the right hand side. There was a significant linear relationship between the two correlation populations (*r* = 0.77, *p* < 10^−6^) indicating that the Post DFC estimates largely retain the correlation with the nuisance norms that is observed in the Pre DFC estimates.

### 3.5. Block regression is similarly ineffective in removing nuisance effects from the DFC estimates

As an alternative nuisance removal approach, we performed block regression in which nuisance measurements were projected out from each window separately. In Figure 8, we superimposed plots of the DFC estimates after performing block regression (green lines; referred to as Post BlockReg DFC) on the plots previously shown in Figure 1 for the BS002 dataset. The correlations between the Post BlockReg DFC estimates and nuisance norms ranged from *r* = 0.63 to *r* = 0.86 with a mean correlation of *r* = 0.70 and per-scan *p* < 0.03. For the CFMRI dataset, the corresponding qualitative figures for block regression results are given in Supplementary Figure 5.

Viewed across the entire sample, the Post BlockReg DFC correlation values shown in Figure 3b ranged from a minimum of *r* = −0.89 to a maximum *r* = 0.87. The correlation distributions were similar to the Post FullReg DFC histograms with cosine similarity values of *S* = 0.93 and *S* = 0.95 for the correlation and absolute correlation values, respectively.

Over the sample, the mean absolute correlation |*r*| between the Post BlockReg DFC estimates and nuisance norms across all scans was significant with *p* < 10^−6^ (see Figure 5a). We also found that 14% of the correlations between the nuisance norms and Post BlockReg DFC estimates were significant at the *p* < 0.05 level and 22% of the correlation values were significant at the *p* < 0.10 level.

Lastly, in Figure 7b, we plot the correlations between the Post BlockReg DFC estimates and the nuisance norms versus the correlations between the Post FullReg DFC estimates and the nuisance norms. A linear fit between the two correlation populations (shown with solid black line) revealed a significant linear relationship (*R*^2^ = 0.86, *r* = 0.93, *p* < 10^−6^). In addition, the correlation distributions for full and block regression were not significantly different from each other (*p* = 0.86, paired two-tailed t-test). The effect size (*d* = 0.0017) and the absolute difference in correlation population means (0.0006) were negligibly small. Thus, with respect to the relationship between the DFC estimates and the nuisance norms, block and full regression have nearly identical effects.

**Figure 8:**
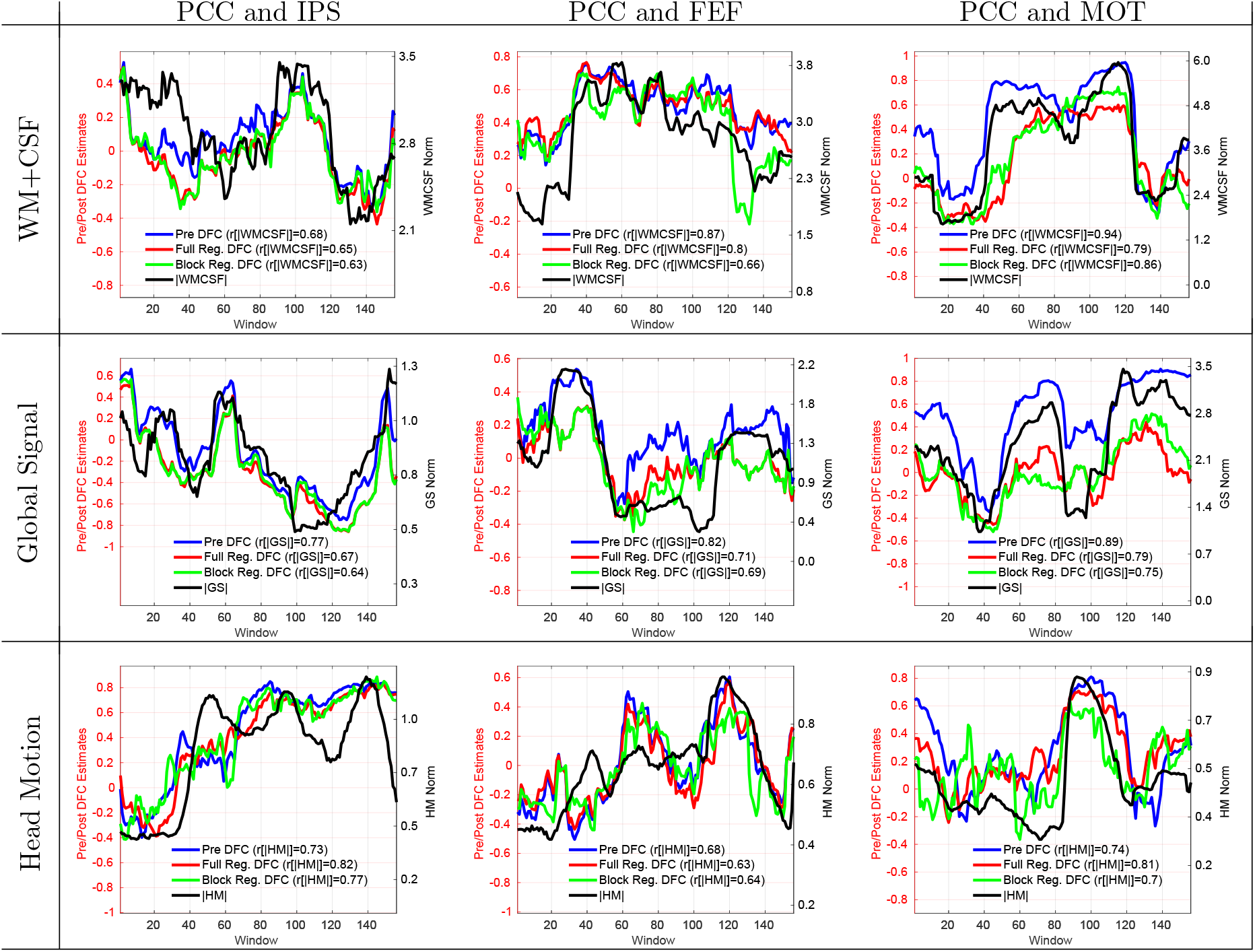
Same 9 representative scans from Figure 1 with the DFC estimates after block regression (solid green line) superimposed. The block regression DFC estimates were significantly correlated (*p* < 0.03) with the respective nuisance norms with a mean correlation value of *r* = 0.70.

In the results presented so far, we have looked at the effects of nuisance regression when using different groups of regressors separately (e.g. WM and CSF, RVf and HRf, GS, and the 6 HM regressors). Note that multiple regression was used when there were more than one regressor in a group. In Supplementary Figures 6 and 7, we show that similar results are obtained when performing multiple regression with (a) WM and CSF grouped with the 6 HM regressors and (b) GS grouped with WM, CSF, and the 6 HM regressors.

## 4. Interpretation

### 4.1. Nuisance effects on correlation estimates

In Section 3.1, we showed that DFC estimates can be related to the norms of various nuisance terms. Here, we aim to provide an intuitive understanding of how this relationship might arise. We use simple toy examples to demonstrate the key principles and to establish concepts that will be further developed in the Theory section. In the toy examples, we represent time series as vectors in a low-dimensional (2D or 3D) space, such that the correlation between time series is simply the cosine of the angle between the vectors. For all of the examples, we will assume that there is a set of two underlying vectors with a fixed angle across time windows, corresponding to an idealized case in which the windowed correlation between two time series is fixed across time. Then, we examine what happens when a nuisance term is added to the underlying vectors. Note that due to considerations of simplicity and mathematical tractability, we restrict our presentation to the case of a single nuisance regressor in both this section and the following Theory section.

### 4.2. 2D Examples

#### 4.2.1. Positive correlations (Aligned Case)

We begin with a simple 2D toy example in which an additive nuisance term points in the same general direction as the underlying vectors. The underlying vectors are depicted with green vectors and denoted as *y*_1_ and *y*_2_ in Figure 9a for 5 consecutive windows (*k* through *k* + 4). The angle between the vectors is fixed at an angle of 135°, such that the correlation is *r* = −0.70 for each window. We consider an additive nuisance term pointing along the horizontal axis with a norm that varies across windows (indicated by the red vectors and denoted as *n_k_* through *n*_*k*+4_).

**Figure 9:**
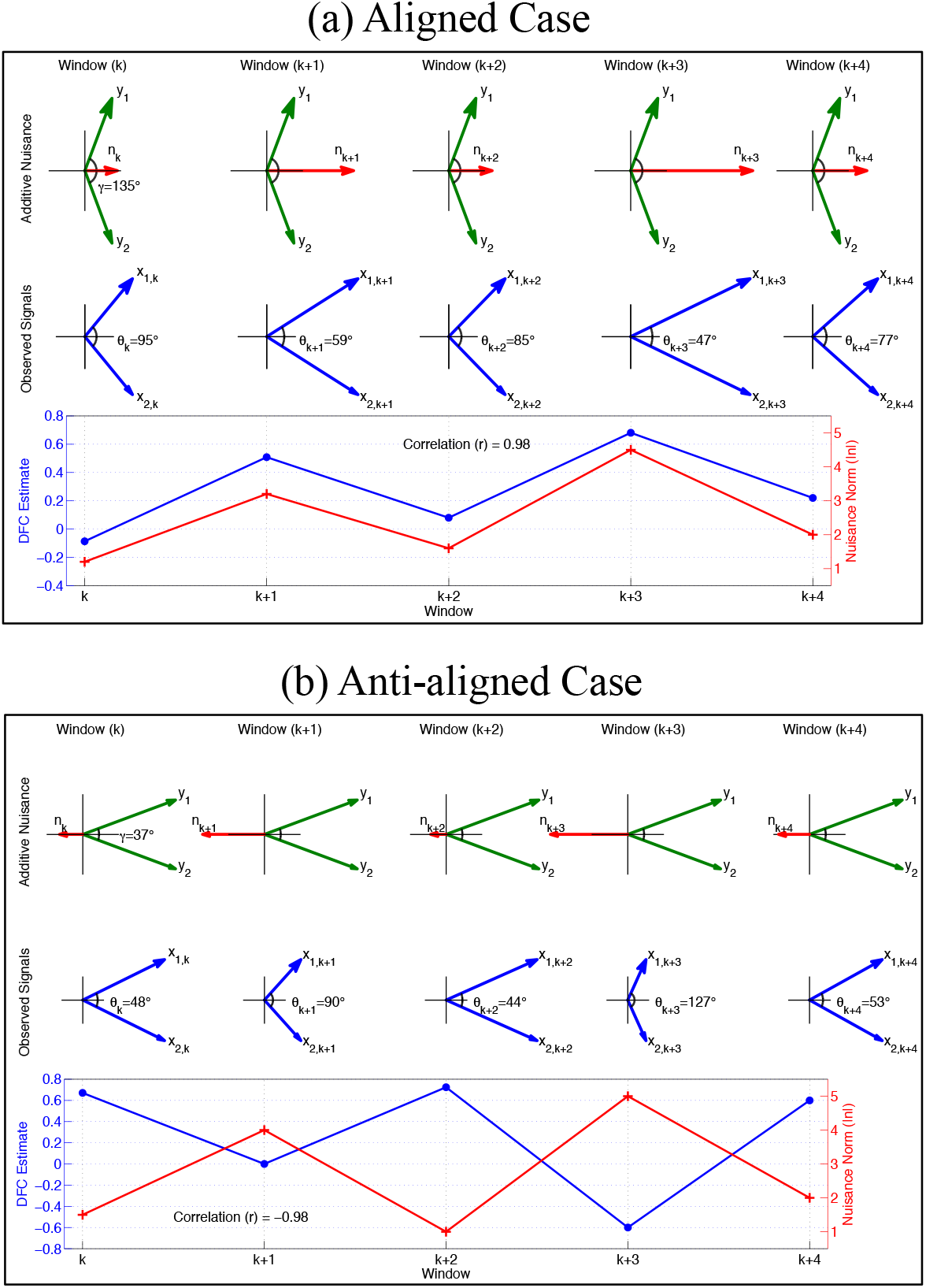
The relationship between the DFC estimates and the nuisance norms under an additive nuisance model. We present two cases indicated as the (a) aligned case and the (b) anti-aligned case. In the first row in (a), we simulate a set of fixed underlying vectors *y*_1_ and *y_2_* across 5 windows (green arrows). The nuisance vectors have different norms in each window (red arrows). The observed time courses *x*_1,*k*_ and *x*_2,*k*_ in the second row (shown with blue arrows) are the sum of the corresponding underlying vectors (*y*_1_ and *y*_2_) and the nuisance vector *n_k_*. In the last row in (a), the simulated DFC estimates (cos*θ_k_*, solid blue line) follows the norm of the nuisance term (|*n_k_*|, solid red line). The aligned case shows that an increase in the norm of a nuisance vector that points in the direction of the underlying signals decreases the inner angle between the observed time courses and increases the value of DFC estimates. The anti-aligned case in (b) shows that an increase in the norm of a nuisance vector, that points in a direction opposite to that of the underlying signals, can decrease the DFC estimate and lead to anti-correlation between the nuisance norm and the DFC estimate.

In this example, the nuisance vectors *n_k_* are aligned with the underlying vectors *y*_1_ and *y*_2_, meaning that the inner products 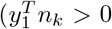 and 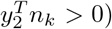 are positive. The sum of the underlying vectors and additive nuisance terms yields the observed time series as shown by the blue vectors in the second row of Figure 9a and denoted as *x*_1,*k*_ and *x*_2,*k*_. The angle between the observed vectors (shown with *θ*_k_) varies with the nuisance norm, with smaller angles observed for larger nuisance norms. In the last row of Figure 9, we plot the correlation between the observed signals (cosine of the angle between observed signal vectors) and the nuisance norm across windows. There is a strong relationship (*r* = 0.98) between the window correlation values and the nuisance norm. This simple example shows how variations in the norm of a nuisance term may induce variations in the windowed correlation estimates that are highly correlated with the norm of the nuisance term.

#### 4.2.2. Anti-correlations (Anti-aligned Case)

As noted in Section 3.1, the DFC estimates can sometimes be anti-correlated with the nuisance norms. To see how this might arise, we consider the ‘anti-aligned’ case shown in Figure 9b. In contrast to the aligned case discussed above, the nuisance vector points in a direction opposite to the average direction of the underlying signals, such that the corresponding dot products 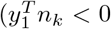 and 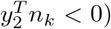 are negative. In this example, the correlation between *y*_1_ and *y*_2_ is fixed at a value of *r* = cos 37° = 0.80.

The addition of the anti-aligned nuisance vector leads to cancellation of vector components along the horizontal axis. As a result, the angle between the observed vectors *x*_1,*k*_ and *x*_2,*k*_ in the second row tends to increase as the nuisance norm increases. Taking the cosine of the angle between the observed vectors to obtain the correlation estimates, we find that the correlation values are anti-correlated (*r* = −0.98) with the nuisance norm, as shown in the third row of Figure 9b. This simple example shows that an additive nuisance term can induce variations in the windowed correlation estimates that are anti-correlated with the nuisance norm.

We should note that if the nuisance norm becomes extremely large for the anti-aligned case, the observed vectors can be dominated by the nuisance term in a manner that can cause the nuisance norm to be positively correlated with the resulting DFC estimates (see the first three windows *n_k_* through *n*_*k*+2_ in Supplementary Figure 8 for an illustration). It is possible that this mechanism may contribute to the positive skew observed in the correlation histograms in Figure 3a-b. However, in the absence of ground truth it is not currently possible to make any definitive conclusions in this regard. In general, the extent to which a nuisance term induces positive or negative correlations between the correlation estimates and the nuisance norm depends on both the angle between the underlying vectors and the relative direction and magnitude of the nuisance vector.

Finally, in Supplementary Figure 8 we present a last 2D example in which the relationship between the DFC estimates and nuisance norms exhibits both positive and negative correlations such that the overall observed correlation is *r* = 0. The nuisance norm is extremely large in the first three windows (*k* through *k* + 2) and dominates over the underlying vectors such that the nuisance norm is positively correlated with the resulting DFC estimates. The nuisance norm then becomes much smaller in the last two windows (*k* + 3 and *k* + 4) so that the nuisance term is no longer able to dominate over the underlying vectors and the nuisance norm is anti-correlated with the resulting DFC estimates. Because of this varying relationship between the DFC estimates and nuisance norms, the observed correlation across all windows is equal to *r* = 0 even though there is a clear effect of the nuisance term on the observed vectors. In general, the relationship between DFC estimates and nuisance norms will exhibit a highly variable and complex behavior across different scans, such that there are many other possible scenarios (in addition to the simple one presented here) that could be used to explain the observation of small correlations between DFC estimates and nuisance norms.

### 4.3. Extension to 3D with addition of an orthogonal nuisance component

In the 2D examples discussed so far, the nuisance term was completely within the 2D plane spanned by the observed vectors. We now expand the example to 3D by including an additional nuisance component that is orthogonal to the 2D plane, such that the overall nuisance term is the sum *n* = *n_I_* + *n_O_* of an in-plane component *n_I_* that lies in the subspace spanned by the observed vectors *x*_1_ and *x_2_* and an orthogonal component *n_O_* that is orthogonal to the subspace.

In Figure 10, we have constructed the observed signals (blue vectors in the first row) using the fixed vectors and in-plane nuisance components previously used in the 2D example of Figure 9a. We then modify the nuisance terms by adding orthogonal components, such that the overall nuisance terms (red vectors) have both in-plane and orthogonal components. We construct the orthogonal component in each window such that its norm is proportional to the norm of the in-plane component (i.e. |*n_O,k_*|∝| *n_I,k_*|). Here, it is useful to define a metric |*n_O_*|^2^/(|*n_I_*|^2^ + |*n_O_*|^2^) = |*n_O_*|^2^/|*n*|^2^ that reflects the relative fraction of nuisance energy that lies in the orthogonal component. The orthogonal nuisance fraction is equal to 1.0 when the nuisance component is completely orthogonal to the observed signal subspace and is equal to 0.0 when the nuisance component lies within the subspace. For this example, the orthogonal nuisance fraction (indicated by the black dots in the fourth row) is relatively large and fairly constant with a mean value of 0.94.

**Figure 10:**
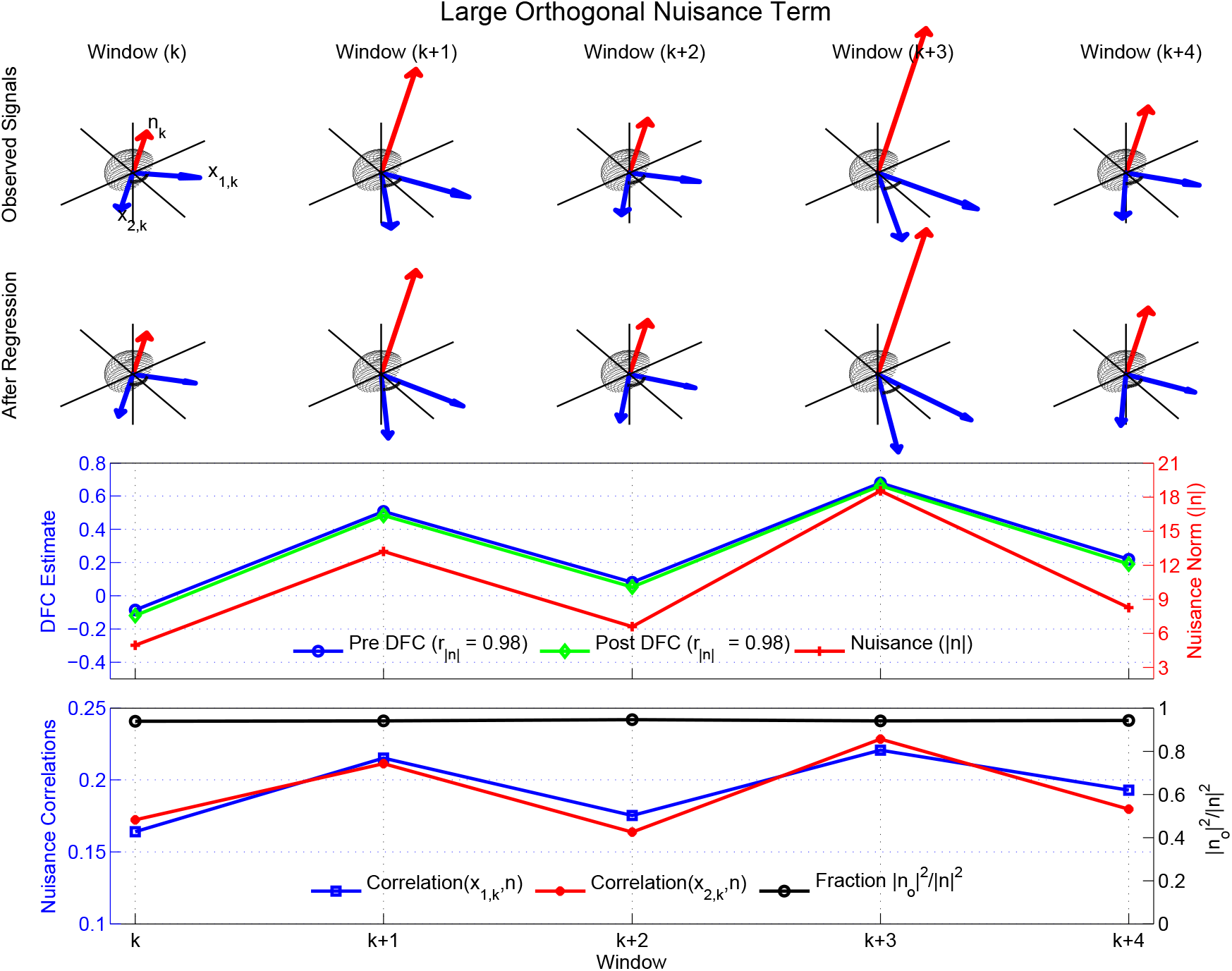
Linear regression can fail to eliminate the relationship between the nuisance norm and the DFC estimates due to a large degree of orthogonality between the nuisance measurement and fMRI time courses. The observed signals (blue vectors) were obtained using nuisance terms (red vectors) for which the in-plane nuisance components are the same as those previously used in Figure 9a. The Pre DFC estimate (blue line) is highly correlated (*r* = 0.98) with the nuisance norm (red line) . The nuisance term also has a large orthogonal nuisance fraction (indicated by the black dots in the fourth row) with a mean value of 0.94. Therefore, linear regression has a minimal effect and Post DFC estimates (green line) are also highly correlated (*r* = 0.98) with the nuisance norm. The correlations between the nuisance term and the observed vectors (red and blue lines in bottom row) are relatively weak. Note that for display purposes, the orthogonal axes (vertical) in the top 2 rows are compressed (×0.7) with respect to the in-plane axes.

The resulting 3D example has several important properties. First, the Pre DFC estimates (blue dots in the third row) exhibit a strong correlation (*r* = 0.98) with the nuisance norm (red dots), similar to that previously shown for the 2D example in Figure 9a. This is because (1) the observed signals are the same in the 2D and 3D examples and (2) by construction the nuisance norm of the 3D nuisance term scales with the norm of the in-plane component.

Second, because the orthogonal nuisance fraction is relatively large, the underlying correlations between the nuisance terms and the observed vectors (i.e. cosine of the angle between the red and blue vectors) are relatively small as shown by the red and blue curves in the fourth row. Thus, this example is consistent with the empirical findings discussed in Section 3.3: a strong correlation between the Pre DFC estimates and the nuisance norms can exist even when the correlation between the nuisance and seed signals is small and the orthogonal fraction is high.

Third, linear regression has a minimal effect in this example, such that the signal vectors after regression (blue vectors in the second row) are similar to the original observed signals (blue vectors in the first row). Note that to simplify the presentation, we have used block regression to obtain the Post DFC estimates. The resultant Post DFC estimates (shown with solid green line and diamond markers in the third row) are also highly correlated (*r* = 0.98) with the nuisance norm. This is consistent with the empirical findings from Sections 3.4 and 3.5 that nuisance regression can have a minimal effect on the relation between the DFC estimates and the nuisance norms.

To aid in relating the orthogonal nuisance fraction to the experimental results, Figure 11 plots the orthogonal fraction values (averaged over each scan) versus the average percent variance in the seed time courses that is explained by the nuisance regressors (computed as the square of the RMS correlation values previously shown in Figure 6). As the average percent variance explained increases, the fraction of the nuisance term energy that is orthogonal to the plane spanned by the observed time series decreases. As noted in Section 3.3, 36% of the strong correlations between the DFC estimates and nuisance norms occurred when the percent variance explained was less than 6.25%. Using the linear fit shown in Figure 11, this percentage corresponds to orthogonal nuisance fractions greater than 0.80, consistent with our use of a large orthogonal fraction in the example.

### 4.4. Regression effects depend on the orthogonal nuisance fraction

In this section, we present a 3D example to demonstrate how the effects of regression depend on the orthogonal nuisance fraction. In each of the 5 cases shown in Figure 12 the nuisance vector n has different orientations with respect to the 2D subspace spanned by the observed signals *x*_1_ and *x*_2_.The orthogonal fraction starts at 1.0 for Case 1 and then decreases to 0.0 for Case 5, with intermediate values for the other cases. The correlation between the observed time courses is fixed at *r* = 0.78 for all of the cases. In the middle column, we show the vectors (denoted as *x̃*_1_ and *x̃*_2_) after nuisance regression.

**Figure 11:**
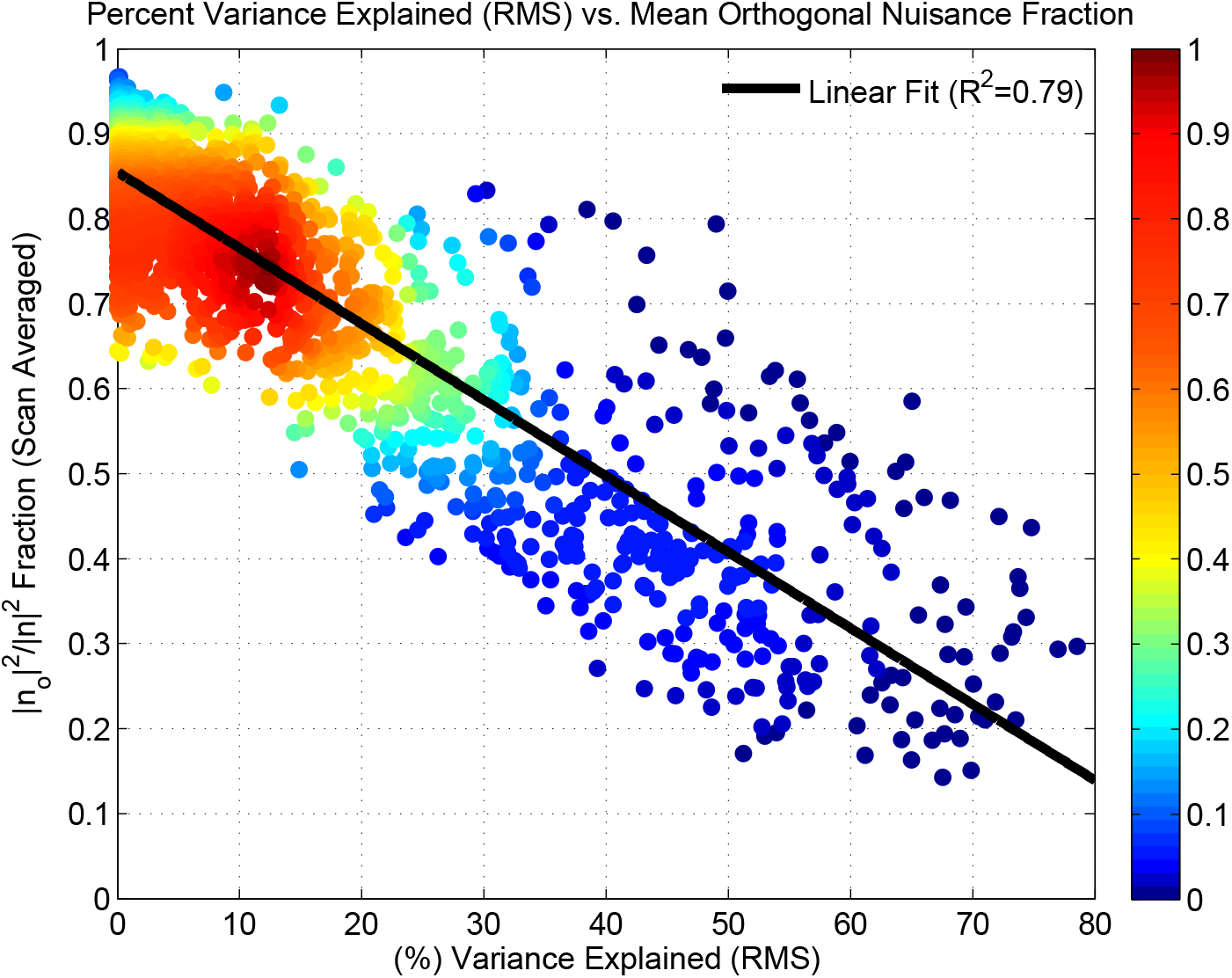
The scan-averaged orthogonal nuisance fraction |*n_o_*|^2^/|*n*|^2^ versus the percent variance explained by the raw nuisance regressors (squared RMS correlation values from Figure 6). As shown by the black least squares line, the orthogonal fraction decreases in a linear fashion (*R*^2^ = 0.79) with increasing percent variance. The relative density of values (maximum density is normalized to 1.0) is indicated by the color map on the right-hand side.

When the orthogonal fraction is 1.0 (Case 1), the nuisance vector is completely orthogonal to the space spanned by the observed signals, and therefore the regression coefficient between the nuisance vector and each observed signal is zero. As a result, linear regression has no effect, and the post regression signals are identical to the observed signals. Consequently, for Case 1 the post regression correlation value is equal to the pre regression correlation value, as shown in the rightmost plot in Figure 12 as the intersection between the red dot and the horizontal green line representing *r* = 0.78.

When the orthogonal fraction is 0.0 (Case 5), the nuisance vector lies completely in the plane spanned by the observed time courses. After regression, the vectors *x̃*_1_ and *x̃*_2_ must point in opposite directions in order to achieve orthogonality to the in-plane nuisance term. As a result, the correlation between the post regression time courses will be −1. (Note that if the nuisance term *n* is in-plane but lies outside of the inner angle formed by *x*_1_ and *x*_2_ then regression will force *x̃*_1_ and *x̃*_2_ to point in the same direction and the correlation will be forced to be +1. In the Appendix, we refer to this as the complementary case).

**Figure 12:**
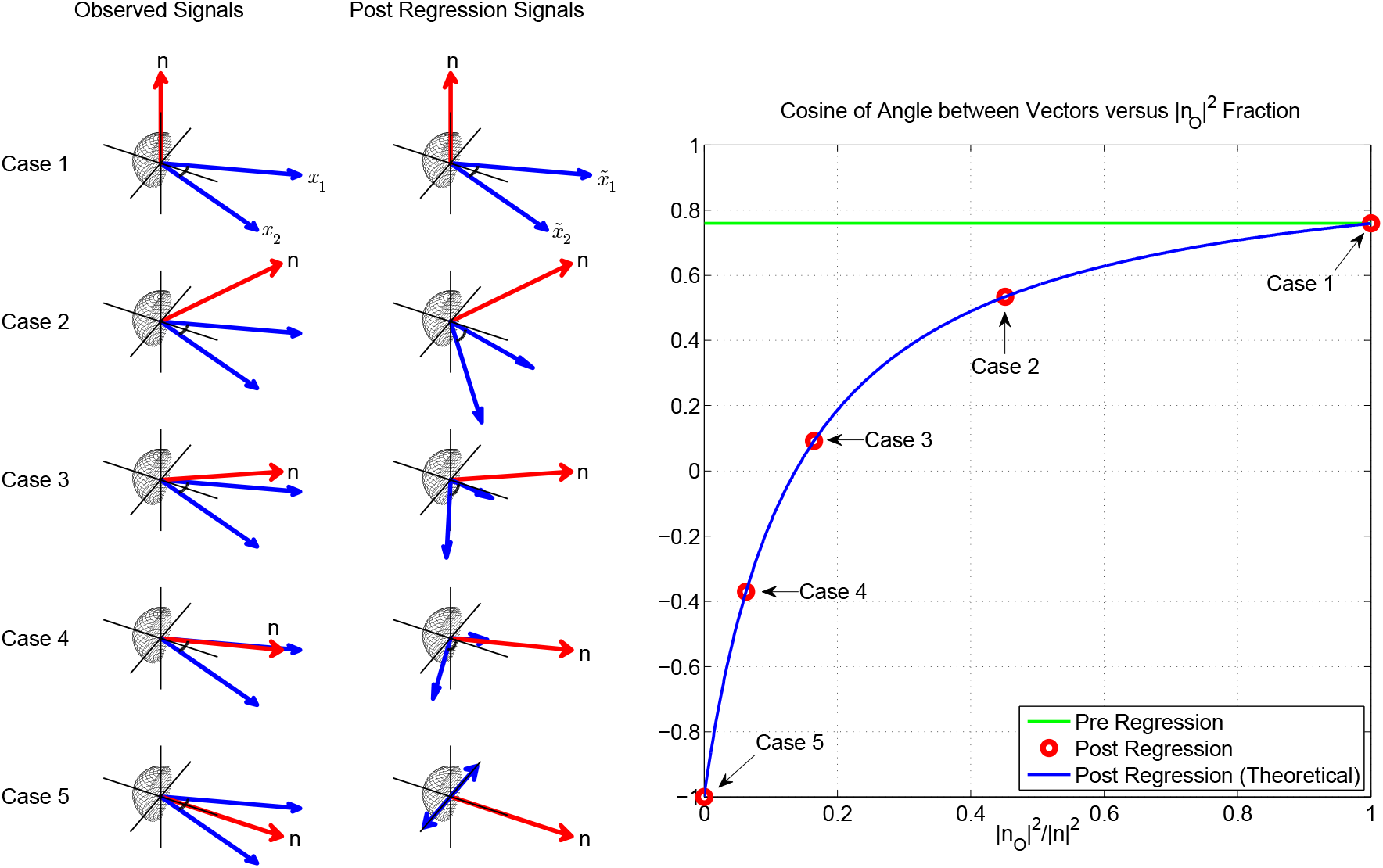
The effect of linear regression on cosine of angle between two vectors. We illustrate 5 cases for the same observed fMRI time courses with a fixed correlation value (*r* = 0.78; as shown with the green solid line) and a variable nuisance component. In Case 1, the nuisance vector is orthogonal to the observed vectors and the orthogonal nuisance fraction (|*n_O_*|^2^/|*n*|^2^) is 1.0, in which case regression has no effect on the pre regression correlation values. Thus, the post regression correlation value is also *r* = 0.78 as shown with the red marker. In Case 5, nuisance vector lies within the observed vectors with |*n_O_* |^2^ / |*n*|^2^ = 0 and post regression correlation value is −1.0 regardless of the pre regression correlation value. For Cases 2, 3, and 4, the fraction |*n_O_* |^2^/|*n*|^2^ takes on intermediate values, such that the difference between the pre and post values increases as the orthogonal fraction decreases. The blue line represents the theoretical values obtained from Equation 2.

For the remaining cases, the post regression correlation values vary as a function of the orthogonal nuisance fraction as shown by the red circles on the right hand side of Figure 12. The green line in the plot indicates the pre-regression signal correlation values (*r* = 0.78), which are independent of the fraction. Cases with a larger orthogonal nuisance fraction have post regression correlation values that tend towards the pre regression correlation value of *r* = 0. 78, whereas cases with a smaller orthogonal nuisance fraction have post regression correlation values that tend towards —1. The blue line in the plot shows the theoretical relation between the post regression correlation values and the orthogonal nuisance fraction. This relation is discussed in greater detail in the Theory section.

Overall, we see that when the orthogonal nuisance fraction is relatively high (e.g. greater than 0.5), the difference between the pre and post regression DFC estimates (distance between blue and green lines) will be relatively small. The exact bound on this difference (over all possible pre-regression correlation values) is provided in the Theory section, where it is also shown that most nuisance regressors lie within the high orthogonal fraction regime and therefore exhibit a small difference between the pre and post regression DFC estimates. The primary exception is the global signal, which provides the motivation for our next example.

### 4.5. DFC estimates after regression with smaller orthogonal nuisance term

In this subsection, we consider a second example in which the orthogonal nuisance fraction is relatively modest with a mean value of 0.47 across windows, as shown with the black line with circles in the fourth row of Figure 13. As discussed further in the Theory section, this value for the orthogonal nuisance fraction is consistent with what is observed for the global signal nuisance term. In this case, linear regression has a noticeable effect as can be seen by the difference between the original signal vectors (blue vectors in the top row) and the post regression vectors (blue vectors in the second row). Regression moves the post regression correlation values away from the pre regression correlation value towards —1.0, similar to Cases 2 and 3 in Figure 12. However, since the orthogonal nuisance fraction is relatively constant across windows, the difference between the Pre and Post DFC estimates is also fairly constant. As shown in the third row in Figure 13 the Post DFC estimates can be approximated as a shifted version of the Pre DFC estimates. Because correlation is invariant to constant offsets, the correlation (*r* = 0.99) between the Post DFC estimates with the nuisance norm is essentially the same as the correlation (*r* = 0.98) between the Pre DFC estimates and the nuisance norm. The construction of the toy example is consistent with our overall observation of a fairly constant difference between the Pre and Post DFC estimates obtained after GS regression. A real example scan with slightly smaller but confined orthogonal nuisance fraction is shown in the second row of Figure 14 for GS regression. In Section 5.3 we provide additional empirical and theoretical results regarding this effect.

**Figure 13:**
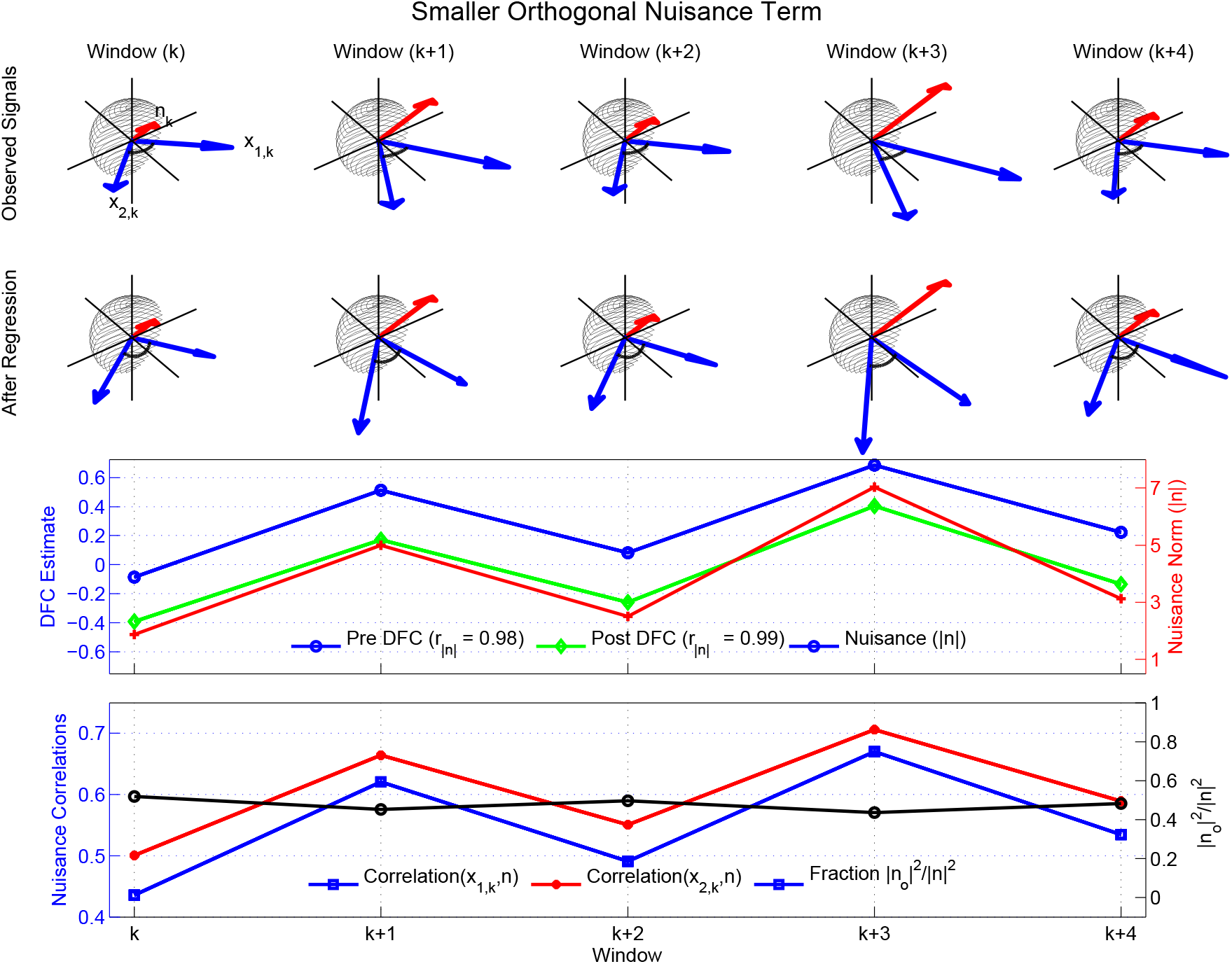
Toy example illustrating how linear regression can fail to eliminate the relationship between the nuisance norms and the DFC estimates when the nuisance orthogonality is weak. Similar to the first toy example in Figure 10, the in-plane nuisance components are the same as those previously used in Figure 9a, however, the orthogonal part of the nuisance term in this example is much smaller as compared to Figure 10. Therefore, linear regression has a noticeable effect on the DFC estimates. Since the orthogonal nuisance fraction is fairly constant across windows, the difference between the Pre and Post DFC estimates is also fairly constant. As the correlation coefficient is invariant to constant offsets, the correlation between the nuisance norm and Pre DFC estimate (*r* = 0.98) is essentially the same as the correlation between the nuisance norm and the Post DFC estimate (*r* = 0.99).

The correlations between the observed signals and the nuisance terms are shown with the blue square and red dotted lines in the fourth row of Figure 13, with values ranging from *r* = 0.44 to *r* = 0.70. Consistent with the smaller orthogonal fraction in this example, these correlations are higher than those observed in the example shown in Figure 10. The range of correlation values used in this example is consistent with the range of RMS correlations that is empirically observed for scans, in which the DFC estimates and GS norm are significantly correlated. Specifically, as shown by the green bars in Figure 6b, the empirical correlations range from *r* = 0.25 to *r* = 0.89 with a mean of *r* = 0.59.

In concluding the Interpretation section, it is important to note that the toy examples considered here are designed to provide a basic level of intuition that can be helpful for understanding both the empirical findings in the Results section and the theoretical expressions which will be presented in the Theory section. While the examples shown here demonstrate behavior similar to that observed in the experimental data, they are by no means exhaustive and alternative examples might be useful to consider in future work.

## 5. Theory

### 5.1. The DFC estimate after block regression

In this section, we provide the expression for the DFC estimate after block regression. The detailed steps of this derivation can be found in Appendix A. Dropping the window index subscript *k* for simplicity, the correlation coefficient after block regression for a single window is given by:

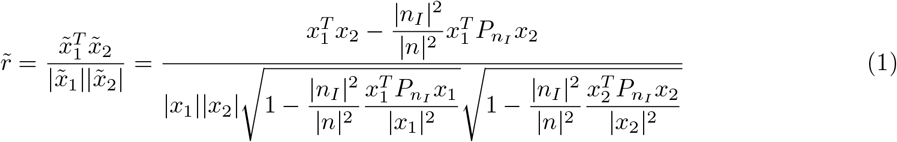

where *x*_1_ and *x*_2_ are the time course column vectors prior to regression, *̃x*_1_ and *̃x*_2_ are the time course column vectors after regression, *n* and *n_I_* are the nuisance term column vector and the in-plane component (also a column vector), respectively, and *P_nI_* denotes the projection matrix onto the in-plane nuisance component.

This can rewritten as

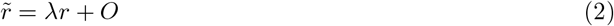

where 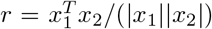 is the correlation coefficient before regression and the scaling λ and offset O terms are defined as

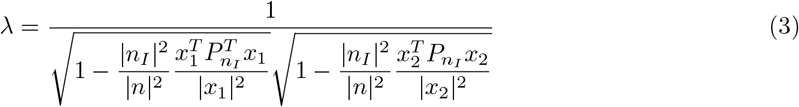

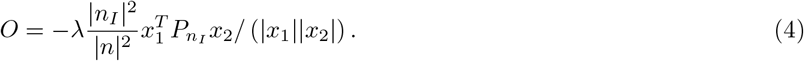

As the orthogonal component *n_O_* becomes arbitrarily large compared to the in-plane component *n_I_*, then the orthogonal fraction 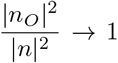 and the term 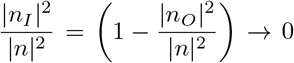 In this case, the scaling term λ > 1 and the offset term *0* > 0, such that the correlation coefficient after regression approaches the pre-regression value *r̃* > *r*. This corresponds to Case 1 in Figure 12.

On the other hand, when the orthogonal component becomes arbitrarily small and the orthogonal nuisance fraction 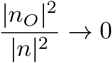, it can be shown that *r̃* approaches either −1.0 or 1.0 (see Equation (A.23) in the Appendix). When the terminal value is −1.0, this corresponds to Case 5 in Figure 12. For intermediate values 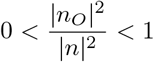 of the orthogonal nuisance fraction, the correlation coefficient *r̃* takes on values between −1 and *r*, corresponding to Cases 2 through 4.

### 5.2. A mathematical bound on the change in DFC using block regression

Here, we consider the difference ΔDFC = *r̃* − *r* between the correlation coefficients obtained before and after block regression. In Appendix B, we show that this quantity is bounded as follows:

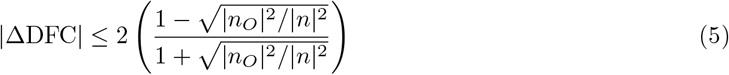

As an example of this bound, in the second column of Figure 14 we plot ΔDFC versus the orthogonal nuisance fraction 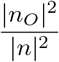 for WM regression (first row) and GS regression (second row) applied separately to two representative scans. Consistent with the discussion in the previous sections, when the orthogonal nuisance fraction 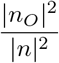 is close to 1.0, the theoretical bound (black dashed line) approaches 0.0. Consequently, the post regression correlation coefficients for the WM signal in the first column of Figure 14 are constrained to be close to the pre regression coefficients.

**Figure 14:**
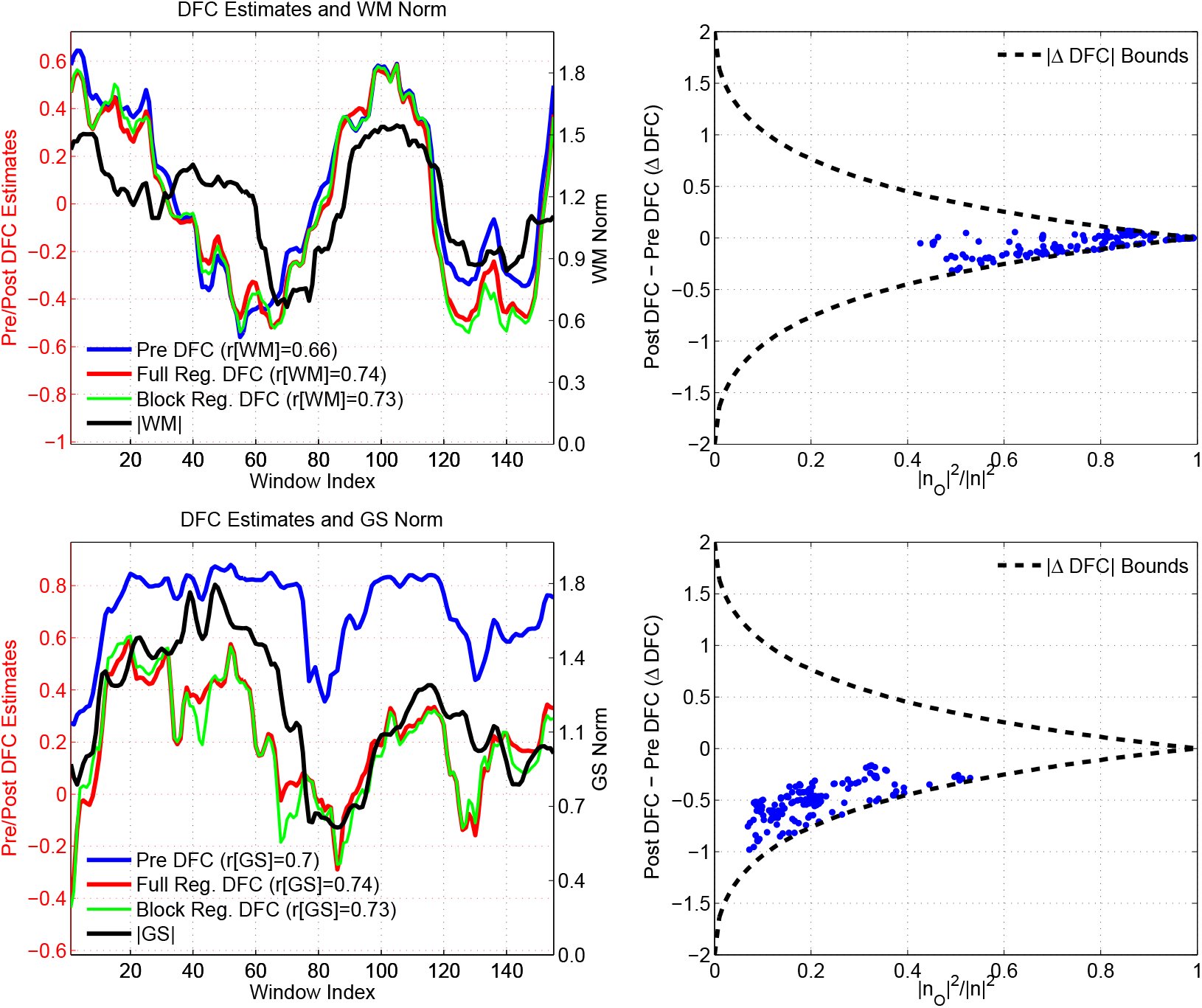
Two representative scans that demonstrate the theoretical bounds on ΔDFC. In the first column, we show the DFC estimates obtained between the PCC and IPS seeds. In these scans the DFC estimates show a large degree of correlation with the norm of the WM time course (*r* = 0.66) in the first row, and with the norm of the GS time course (*r* = 0.7) in the second row. Performing full or block regression does not reduce these correlations and the correlations are larger than 0.7 after each regression technique. In the second column, we show the ΔDFC values versus the orthogonal nuisance fraction for the WM and GS regressors. Each point in these plots corresponds to the orthogonal nuisance fraction and ΔDFC values in a specific window. We also superimpose the theoretical bounds. For the WM nuisance term, the effects of regression are limited by the tight bound imposed by the large orthogonal nuisance fractions. For the GS nuisance term, the bounds are more relaxed due to the smaller orthogonal nuisance fraction. However, the ΔDFC values are clustered around −0.5 and the difference between the Pre DFC and Post DFC values in the first column is also fairly constant. Thus, the Post DFC values remain correlated with the nuisance norms after GS regression.

On the other hand, as the orthogonal nuisance fraction 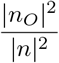 approaches zero, the theoretical bound relaxes and approaches ±2. This corresponds to the case where the post regression correlation coefficient approaches either −1.0 or 1.0. Since the pre regression correlation coefficient is bounded between −1.0 and 1.0, the maximum absolute difference in coefficients is 2, consistent with the theoretical bound on ļΔDFC|. The post regression correlation coefficients for the GS signal in the first column of Figure 14 have a noticeable negative offset when compared to the pre regression coefficients.

To demonstrate the general validity of the bound, in Figure 15, we plot ADFC versus the fraction 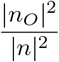 for six different nuisance regressors using the data from all scans and different seed pairs. For head motion (HM), the nuisance regressor was defined as the first principal component of the 6 motion regressors. All of the empirical DFC differences are found to lie within the theoretical bounds. For WM, CSF, HM, RVf, and HRf regressors, the mean of the orthogonal fractions ranged between 0.78 and 0.83, reflecting the fact that for most of the data windows there is a large orthogonal fraction and a fairly tight bound on ΔDFC such that mean difference between the pre and post regression values is small, ranging from −0.049 to −0.0065. In contrast, for the GS regressor, the mean orthogonal fraction is 0.47, and so the data for most of the windows lie in a range where the bounds on ADFC are not as tight and the mean difference between pre and post regression values is −0.26. The smaller orthogonal nuisance fraction observed for GS reflects the fact that it is derived as the mean of all the voxel time courses in the brain and therefore is expected to exhibit a greater similarity (and hence a greater in-plane component) with the seed voxel time courses, as compared to the other regressors.

**Figure 15:**
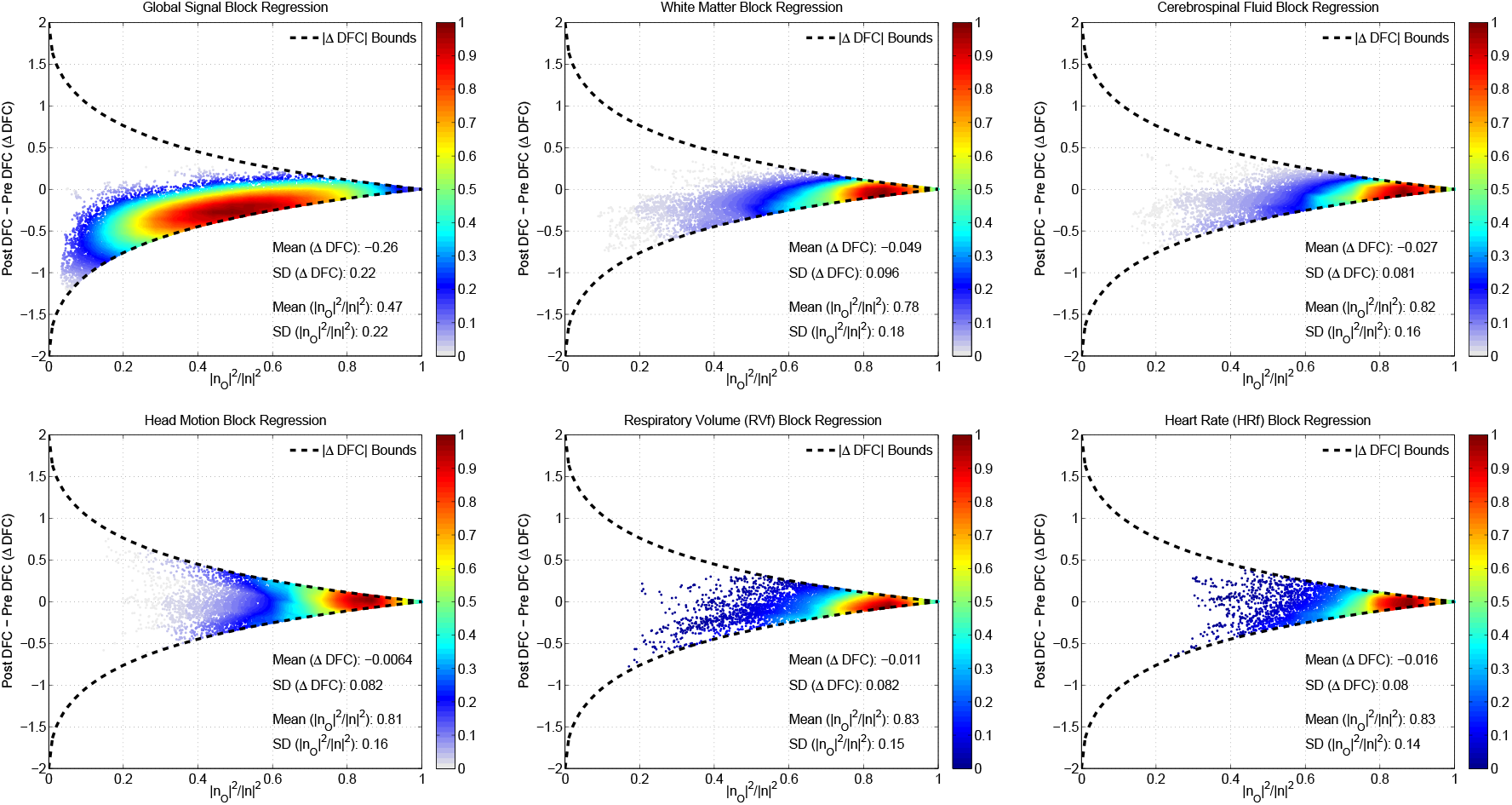
ΔDFC versus the fraction 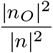 for six different nuisance regressors (from left to right and top to bottom: GS, WM, CSF, HM, RVf, and HRf) using the data from all scans. Each point in a given plot corresponds to a single window. The theoretical bounds for block regression are shown with the dashed black lines. The empirical ΔDFC values were found to lie within the theoretical bounds. For WM, CSF, HM, RVf and HRf regressors, the mean of the orthogonal fractions ranged between 0.78 and 0.83. In this region, the bounds are fairly narrow such that the mean differences between the pre and post regression values were small, ranging from −0.049 to −0.0065. For GS, the mean orthogonal fraction was 0.47 and the bounds on ADFC were not as tight. However, the the ΔDFC values were clustered about a mean value of −0.26 and closely followed the lower bound. Note that the relative density of data points (maximum density is normalized to 1.0) is indicated by the color map on the right-hand side of each plot.

### 5.3. Approximate Constant Offset observed in ΔDFC for GS regression

In both the previous section and Section 4.5, we noted that GS regression resulted in a relatively constant difference ΔDFC between the Pre and Post DFC values. In addition, we can see from the upper righthand panel in Figure 15 that the per-window ΔDFC values with GS regression are clustered close to the lower bound.

To further investigate this effect, we computed the mean and standard deviation of the ΔDFC values for each scan. We also computed the mean lower theoretical bound for each scan by computing the lower bound for each window and then averaging over windows. Figure 16a plots the mean ΔDFC values versus the mean theoretical bounds, with the data for GS and non-GS regressors indicated by the circle and square markers, respectively. The line of unity is indicated by the dashed green line. Due to the additive property of inequality, the mean bound is strict so that all points must lie above this line. A linear fit to the GS data is shown with the magenta line (*R*^2^ = 0.79, Slope = 0.78), which is fairly close to the line of unity. This indicates that the mean ΔDFC values after GS regression approximately follow the mean theoretical bound over a large range of ΔDFC values. In contrast, the mean ΔDFC values shown for the non-GS regressors are relatively small, consistent with the large orthogonal nuisance fractions observed for these regressors in Figure 15.

In Figure 16b, we plot the negative standard deviation (NSD) of ΔDFC for each scan versus the average bound, where the standard deviation is negated for display purposes. Note that in contrast to the mean ΔDFC values, there is no requirement that these points lie above the line of unity (dashed green line). The slope of the linear fit for GS (Slope = 0.2, magenta line) is much smaller than the slope of 0.79 observed for mean ΔDFC in Figure 16a. In addition, the magnitudes of the SD values are significantly smaller than the magnitudes of the mean ΔDFC values (p < 10^−6^, paired t-test). Thus, the primary effect of GS regression is to induce a negative offset in the mean of the DFC estimates accompanied by a relatively smaller change in the DFC fluctuations about the mean. As a result, the Post DFC estimates after GS regression can be approximated to first order as a shifted version of the Pre DFC estimates and will therefore largely retain the correlation with the nuisance norm. An example of this limitation was previously shown in the GSR example in the second row in Figure 14.

**Figure 16:**
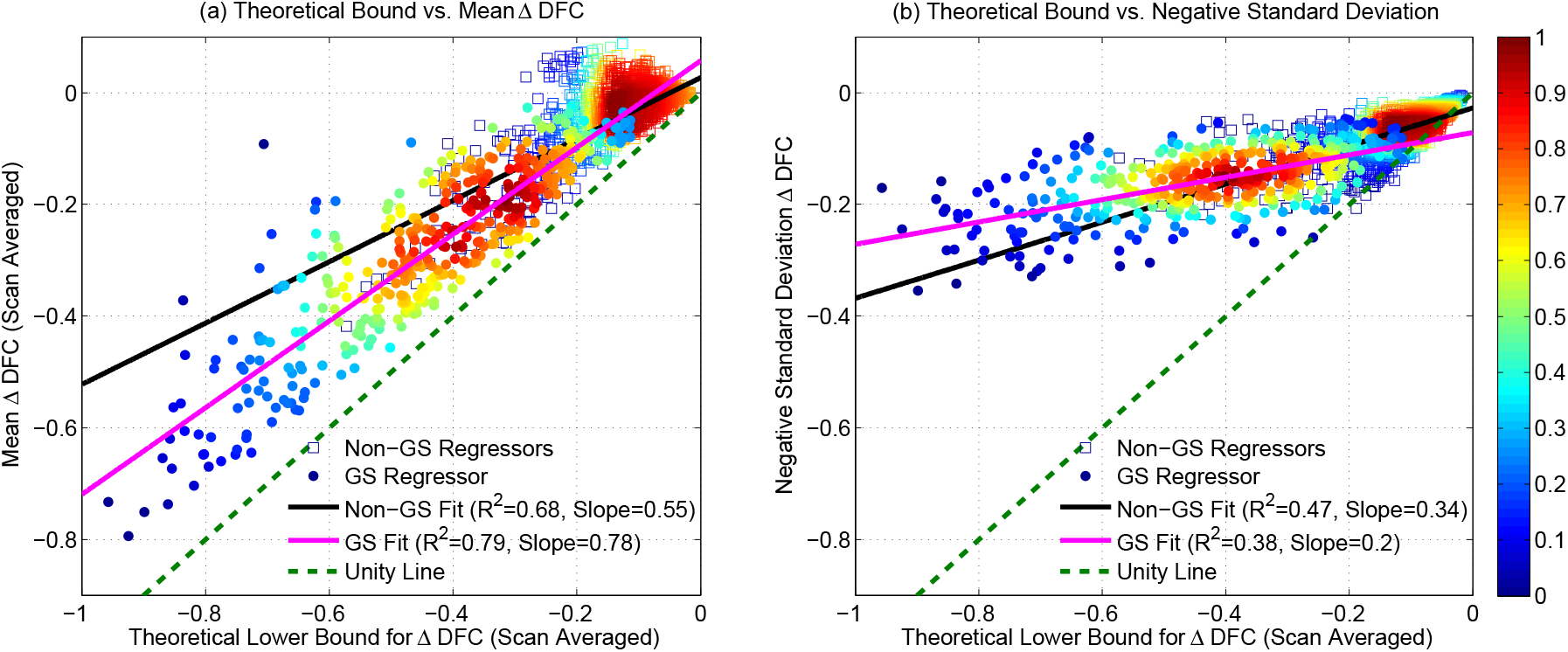
(a) Mean ADFC versus scan-averaged theoretical bound. Data for GS and non-GS regressors are indicated by the circle and square markers, respectively. For GS regression, the linear fit (magenta line, *R*^2^ = 0.79, Slope = 0.78) is close to the line of unity (dashed green line). For non-GS regressors, the linear fit is indicated by the black line and most values occur at small ADFC values. (b) Negative Standard Deviation (NSD) of ADFC (over each scan) versus the mean theoretical bound. For GS regression, both the slope of the best fit line (magenta line, *R*^2^ = 0.38, Slope = 0.2) and the magnitude of the NSD values are considerably lower than the slope and magnitudes observed for the mean ADFC values. The relative density of data points (maximum density is normalized to 1.0) is computed for GS and non-GS data separately and indicated by the color map on the right-hand side.

### 5.4. DFC estimates after full regression

In the Results section we have shown that effects of block regression and full regression on the DFC estimates were very similar. Here, we present the expression for the DFC estimate after full regression as a modified version of the expression obtained for block regression. The main difference between the two approaches is that the regression fit coefficients for full regression are computed from the entire time series data, whereas the fit coefficients for block regression are computed using only the data in the window of interest. However, it important to note that both approaches subtract out a scaled version of the nuisance term in the window of interest.

As derived in Appendix C, the correlation coefficient after full regression for the kth window can be written as:

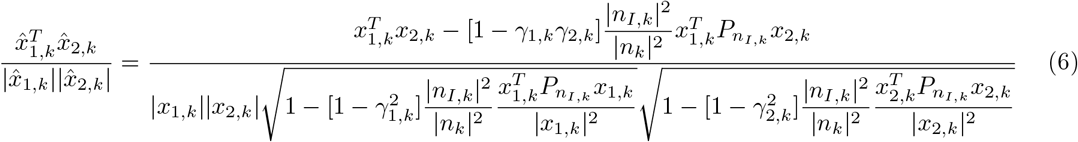

where the hat notation is used to refer to time courses after full regression, and *γ*_1,*k*_ and *γ*_2,*k*_ are scalar correction terms that account for the difference between full and block regression. When these correction terms are zero, the expressions for block and full regression are identical.

The correction terms *γ*_1,*k*_ and *γ*_2,*j*_¾ are empirically determined and vary with the specific features of the signal and nuisance time series. Thus, in contrast to the block regression, it is challenging to derive theoretical bounds on ΔDFC for the full regression case. However, as shown in Figure 17, the empirical values of ΔDFC for full regression are largely within the theoretical bounds obtained for block regression. This is consistent with the empirical similarity of the DFC estimates obtained after block and full regression.

The correction term for a time course *x*_1_ is given in Appendix C as 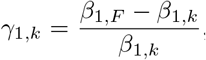, where *β*_1,*F*_ is the regression fit coefficient using full regression and *β*_1,*k*_ is the per-window fit coefficient. As defined, the correction term *γ*_1,*k*_ blows up for very small *β*_1,*k*_ values. However, in practice this term is actually multiplied by *β*_1,*k*_ (see Equation (C.5)), so that it is sufficient to consider the difference between the fit coefficients *m*_1,*k*_ = *β*_1,*F*_ − *β*_1,*k*_. Similarly, for the second time series *x*_2_, we may consider the difference term *m*_1,*k*_ *m*_2,*k*_. Note that as *m*_1,*k*_, *m*_2,*k*_, and the product term *m*_1,*k*_*m*_2,*k*_ go to zero, then *γ*_1,*k*_, *γ*_2,*k*_, and the product *γ*_1,*k*_*γ*_2,*k*_ all approach zero, and full regression and block regression become identical operations. To provide examples of the behavior of these differences, Supplementary Figure 9 shows that *m*_1*k*_ and *m*_1*k*_ as well as the product *m*_1*k*_*m*_2*k*_ are centered around zero for the GS and WM regressors.

**Figure 17:**
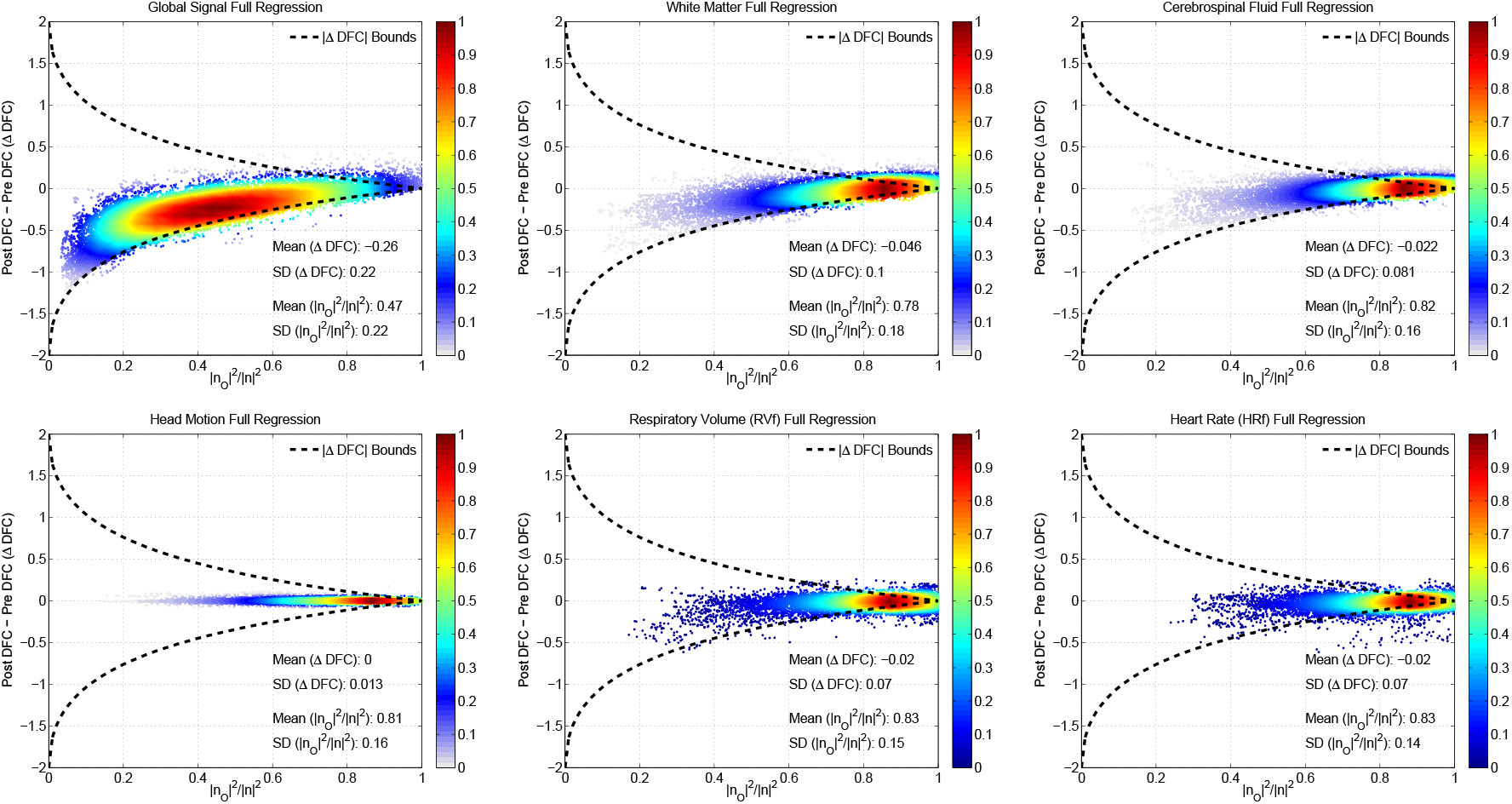
The empirical ΔDFC versus the 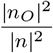 fraction for the same six nuisance regressors used in Figure 15. This | n| ^2^ time full regression was performed to obtain the empirical values but the same bounds are taken from the block regression derivation. The empirical values of ΔDFC for full regression are largely within the theoretical bounds obtained for block regression, consistent with the empirical similarity of the DFC estimates obtained after block and full regression. Note that the relative density of data points (maximum density is normalized to 1.0) is indicated by the color map on the right-hand side of each plot.

## 6. Discussion

### 6.1. Summary

We have shown that sliding window correlation DFC estimates can be strongly and significantly correlated with the sliding window norms of various nuisance measurements. This relationship between the DFC estimates and the nuisance norms can exist even when the correlations between the underlying nuisance and seed time courses are relatively weak. Moreover, we found that significant correlations between the DFC estimates and nuisance norms can persist even after performing nuisance regression. We derived mathematical expressions to describe the effects and limitations of nuisance regression on DFC estimates and demonstrated that the empirical results lie within the theoretically predicted bounds.

Based on our empirical and theoretical findings, we identified two main mechanisms for the inefficacy of nuisance regression. First, as shown in Figure 6, the DFC estimates can be strongly correlated with the nuisance norms even when the underlying correlation between the nuisance terms and the seed signals is relatively low. As a result, for most cases a large fraction of the nuisance term is orthogonal to the subspace spanned by the seed signals. This greatly reduces the efficacy of nuisance regression, such that the difference between the Pre and Post DFC values is relatively small and the relation between the DFC estimates and nuisance norms is largely unaffected. We observed this major limitation of nuisance regression particularly for WM, CSF, RVf, HRf, and HM regressors as demonstrated in Figure 15.

The second mechanism applies primarily to the GS nuisance term which has a smaller orthogonal nuisance fraction. The reduced orthogonal nuisance fraction reflects the fact that the GS is computed as the average of the BOLD time courses across the brain and will therefore tend to have a higher correlation with the seed time courses. As shown in Figure 16, the mean ΔDFC values observed for GS regression closely follow the mean theoretical bound, which approaches −1 as the orthogonal fraction approaches zero. In contrast, the standard deviations of the ΔDFC values exhibit a much weaker dependence on the orthogonal fraction. As a result, the Post DFC estimates after GS regression can be approximated to first order as a shifted version of the Pre DFC estimates, as illustrated in the second row of Figure 14 for a representative subject. Because the Post DFC estimates largely retain the fluctuations in the Pre DFC estimates, they will also retain the correlation with the nuisance norm. Although this mechanism is most often observed for the GS nuisance term, it can sometimes be observed for WM and CSF nuisance terms (e.g. upper righthand panel in Figure 8) in which partial volume effects with gray matter can make the WM and CSF nuisance terms to behave more like the GS.

### 6.2. Nuisance effects in DFC studies

It has been previously noted that nuisance effects might account for a significant portion of the fluctuations in DFC estimates (Preti et al., 2017; Hutchison et al., 2013). The effects may be especially pronounced for sliding window DFC estimates since transient nuisance effects can greatly alter the correlation estimates within a short temporal window (Hutchison et al., 2013). Chang and Glover (2010) looked at the inter-subject correlation between measures of motion and DFC variability, but did not find a significant relation. They also found that intra-subject correlations between sequences of motion-based and DFCs-based deviations were not significant. However, they did not consider the intra-subject correlation between DFC estimates and motion nuisance norms, as was done in this study.

Nikolaou et al. (2016) found that dynamic fluctuations in network degree were significantly correlated with the spectral power of end-tidal CO_2_ and heart rate measurements. Since the network degree calculated in that study was based on the summation of DFC magnitudes, these findings are roughly consistent with our observation of a significant correlation between the DFC estimates and the norms of the RVf and HRf regressors.

Laumann et al. (2017) used multivariate kurtosis to assess deviations from constant covariance and found that the removal of high-motion frames reduced the observed kurtosis. They concluded that a significant portion of the resting state functional connectivity dynamics could be attributable to subject motion.

Glomb et al. (2017) reported that measures of instantaneous BOLD variance (averaged over the entire brain) were significantly correlated with instantaneous average correlation (over all pairwise correlations). These results are consistent with our findings of significant correlations between DFC estimates and the GS norm. A distinction is that the GS norm is the magnitude of an average signal whereas the instantaneous BOLD variance was calculated as the average across variances from different regions. However, prior work has shown that these two measures are highly related (He and Liu, 2012; Wong et al., 2012).

A key finding of this paper is that DFC estimates can be significantly correlated with the nuisance norms even when the underlying nuisance time courses are largely orthogonal to the observed seed time courses. As discussed in the Interpretation section, one plausible scenario that can give rise to this effect is the presence of an additive nuisance term consisting of time-varying in-plane and orthogonal terms, where the magnitudes of the two terms are roughly in sync and the magnitude of the orthogonal term is much larger than that of the in-plane term. Under this scenario, the time-varying in-plane component can cause DFC fluctuations in the observed vectors (since it affects the inner-angle between the observed vectors) even when the correlation of the putative underlying signal vectors (which are not observed) remains constant. Because of the assumed relation between the in-plane and orthogonal terms, the resulting DFC fluctuations are correlated with the overall nuisance norm. In addition, because the in-plane term is much smaller than the orthogonal term, the overall nuisance term exhibits a weak correlation with the observed vectors. It is important to note that while this plausible scenario provides some insight into the empirical observations and helps to motivate the theoretical findings, it is by no means intended to serve as a “model” of the data. The modeling of nuisance effects in fMRI is still an area of active investigation (Liu, 2016; Murphy et al., 2013) and future work will be needed to more fully characterize the impact of nuisance terms on DFC estimates. Such efforts are likely to require a consideration of the non-linear and non-stationary aspects of the underlying signals (Sugihara et al., 2012; Hutchison et al., 2013).

### 6.3. Efficacy of Nuisance regression in DFC studies

Although nuisance regression is widely performed in DFC studies (Hutchison et al., 2013; Preti et al., 2017), its effects on DFC estimates have received relatively little attention. Nikolaou et al. (2016) reported that regression reduced but did not eliminate the correlation between dynamic measures of network degree and measures of heart rate and respiratory spectral power. These findings are aligned with our results showing that regression has a limited effect on the correlation between DFC estimates and HRf and RVf nuisance norms.

In recent work, Xu et al. (2018) found that global signal regression (GSR) had a maximal impact on DFC estimates in temporal windows where the GS mean absolute magnitude was large and a lesser impact in windows where the magnitude was small. They interpreted their findings using the framework introduced in (Nalci et al., 2017b), where it was shown that GSR can be approximated as a temporal down-weighting process. The authors noted that the main effect of GSR was a spatially heterogenous negative shift in the sliding window correlation values, an observation consistent with the predominantly negative ΔDFC values found in this study (see the upper righthand panel in Figure 15). However, in contrast to our study, the authors did not consider the relationship between the DFC estimates and the GS norm.

There is related work for static FC studies regarding the inefficacy of nuisance regression (Power et al., 2017; Chang et al., 2009; Power et al., 2012, 2014). For instance, Power et al. (2017) observed that frame-wise displacement (FD), a summary measure for head motion, remained highly related with resting-state fMRI time courses even after HM regression. As we have discussed in detail, the efficacy of nuisance regression increases as the orthogonal nuisance fraction decreases. Due to the inverse relationship between the orthogonal fraction and the percent variance explained (see Figure 11), this means that the efficacy of regression will decrease as the amount of variance that can be explained by the nuisance regressors decreases. Chang et al. (2009) found that RVf and HRf regressors together explained only 15.8% of the average total variance in voxels which showed a significant correlation between the BOLD time series and nuisance regressors, and the RVf regressor explained only 11 .7% of the average total variance when used on its own. This is roughly in-line with the results shown in Figure 6 where the percent variance values observed for non-GS regressors were largely below 20%, with 54% of the strong correlations between the DFC estimates and non-GS nuisance norms occurring when the percent variance was less than 6.25%.

### 6.4. Other Approaches

In this work, we used the sliding window correlation approach, which is widely used in DFC studies (Preti et al., 2017; Hutchison et al., 2013; Calhoun et al., 2014) both as a primary analysis approach and as an intermediate analysis step (e.g. used to generate DFC estimates that are then further analyzed with k-means clustering or principal components analysis). Other approaches include time-frequency methods such as wavelet transform coherence (Chang and Glover, 2010; Yaesoubi et al., 2015) and probabilistic methods such as hidden Markov modeling (Vidaurre et al., 2017). Regardless of the approach, nuisance regression is a standard preprocessing step using nuisance regressors based on either independent measures (e.g. physiological measurements) or data-driven measures (e.g. GS, WM+CSF, or independent component analysis (ICA) components). Our findings regarding the limited efficacy of nuisance regression for sliding-window DFC estimates suggest that caution must also be exercised when interpreting DFC estimates obtained with other approaches. Nevertheless, future work to assess the impact of nuisance terms and the efficacy of nuisance regression when applied to additional DFC approaches would be of great interest.

In this study, we have examined the use of both physiological (respiratory and cardiac) and data-driven (motion, GS, and WM+CSF) nuisance regressors, which are all widely used in the field (Ciric et al., 2017). There has also been growing interest in the use of ICA-based approaches for denoising of fMRI data. Recent work has pointed out that both traditional nuisance regressor (excluding GS) and ICA-based approaches are limited in their ability to eliminate spatially widespread effects and suggest the continued need for some type of global signal regression (Ciric et al., 2017; Power et al., 2017; Burgess et al., 2016; Power et al., 2018). In the case of ICA-based approaches, the limitations most likely reflect the spatial independence criteria that is inherent to the ICA algorithm. Given this limitation it is unlikely that ICA-based approaches alone can eliminate the relation between DFC estimates and nuisance norms. Nevertheless, the development and comparison of nuisance regressor and ICA-based approaches continues to be an active area of research, and future work to assess the efficacy of ICA-based approaches with regards to the relation between DFC estimates and nuisance norms would be of great interest. In addition, as methods for the removal of nuisance terms continue to evolve, it will be useful to evaluate whether these future approaches can better attenuate the relation between DFC estimates and nuisance norms.

For our examination of the relation between DFC estimates and nuisance norms, we used a seed-based approach in which the average time series from four different seed regions were used. The utilization of the seed based approach with two independent datasets serves as a solid approach for an initial and detailed characterization of the relation between DFC estimates and nuisance norms and the limited efficacy of nuisance regression. An extension of the current work to further characterize these effects using other brain regions and networks would be of interest. This could entail the use of cortical parcellations or ICA-based network components as described in (Preti et al., 2017; Hutchison et al., 2013; Calhoun et al., 2014; Laumann et al., 2017). The fact that the various potential approaches all utilize some type of spatial weighting to derive region-based or network-based time courses suggests that they may yield similar results with regards to DFC estimates and nuisance norms. Nonetheless, further work in this area would be useful to determine if there are any substantial differences.

### 6.5. Nuisance Norm Regression

Given the inefficacy of nuisance regression in reducing the correlations between the DFC estimates and the nuisance norms, it is reasonable to consider alternative approaches. A potential solution is to compute the sliding window norm for each regressor and then project these out from the DFC estimates using linear regression. This procedure is described in Appendix D and referred to as *nuisance norm regression* (NNR). This approach differs from traditional nuisance regression techniques which regress out nuisance measures directly from the fMRI time courses. Instead, NNR acts on the correlation coefficients. Further work is needed to characterize the potential advantages and disadvantages of this approach.

### 6.6. Vigilance Effects

There is a growing appreciation that variations in vigilance may account for a considerable portion of the dynamic fluctuations in resting-state fMRI data (Chang et al., 2016; Falahpour et al., 2018; Haimovici et al., 2017). In addition, the state of vigilance can have an effect on physiological measures, such as the recently reported dependence of pulse oximetry amplitudes on the level of wakefulness (Chang et al., 2018). Our preliminary analysis in Section 3.1 did not reveal an effect of the state of the eyes (open or closed) on the correlations between the DFC estimates and nuisance norms. Nevertheless, future work to assess how the relation between DFC estimates and nuisance norms might depend on the state of vigilance would be of great interest.

### 6.7. Implications for static FC estimates

In this paper, we have considered the relation between nuisance norms and FC estimates obtained across different temporal windows. In DFC studies, the temporal windows under consideration are typically on the order of 40 to 100 second windows. However, the framework for interpreting the observed effects is applicable to arbitrary window lengths. It is therefore of interest to consider whether similar effects are observed for static FC estimates where the window length is equal to the scan duration. In this case both the static FC estimates and nuisance norms are computed over the full duration of each scan and we look to see whether static FC estimates and nuisance norms are related across scans.

In Supplementary Figure 10, we present a preliminary analysis showing the existence of significant correlations (both before and after nuisance regressions) between static FC estimates and nuisance norms across the 68 scans in the BS002 dataset. These preliminary findings are roughly consistent with recent work looking at the correlation between static FC measures and motion-related metrics (Ciric et al., 2017; Siegel et al., 2017; Power et al., 2015). In the preliminary example, the relation between static FC estimates and nuisance norms is examined using different combinations of regressors, as detailed in the figure caption. Similar to the prior work, the inclusion of additional regressors led to a greater reduction (with nuisance regression) in the observed correlations, as compared to the use of only motion regressors. This preliminary work suggests that the framework we have presented may be useful for providing deeper insights into the effects of nuisance terms and regression in static FC studies. Further work that considers the proposed framework within the context of the growing body of related studies on the effects of nuisance terms in static FC studies would be of interest.

### 6.8. Conclusion

We have provided a detailed examination of nuisance effects and regression in DFC measures. Our findings both confirm and significantly extend the limited prior work in this area. In particular, we have shown that DFC estimates can be strongly correlated with nuisance norms even when there is only a weak correlation between the nuisance and seed signals. We demonstrated that nuisance regression is largely ineffective for the removal of this relationship. Furthermore, we provided a mathematical framework to describe the effects of nuisance regression and showed that the experimental findings are in agreement with the theoretical predictions. The current mathematical framework considers a single nuisance regressor and provides valuable insights into the experimental findings. It can be used to approximate the multiple regressor case by taking the first principal component of the regressors, as was done for the HM regressors in Figure 15. While our preliminary efforts indicate that an extension of the framework to readily handle multiple regressors is not straightforward, future work in this area would be of interest.

It is important to note that there is an ongoing discussion in the literature regarding whether DFC estimates are largely due to sampling variability versus true changes in FC (e.g. fluctuations in wakefulness) (Pannunzi et al., 2017; Tagliazucchi and Laufs, 2014; Laumann et al., 2017; Haimovici et al., 2017; Liégeois et al., 2017). The present work does not make any claims as to whether DFC estimates reflect dynamics changes in brain functional connectivity as opposed to sampling variability or other artifactual factors. Instead, the focus is on presenting an empirical finding of a relation between DFC estimates and nuisance norms. This represents an additional factor that should be considered in the interpretation of DFC estimates.

This work highlights a potential confound in the interpretation of DFC studies, which typically make the implicit assumption that nuisance effects are largely minimized in the pre-processing stage. Our findings suggest that the interpretation of DFC measures should be expanded to consider potential correlations with the nuisance norms. If these correlations are not adequately considered, differences in DFC estimates (e.g. between groups or treatment conditions) may be incorrectly interpreted as reflecting meaningful effects when in fact they may be largely attributable to differences in nuisance activity. Because nuisance effects such as subject motion, respiration, and cardiac activity originate in brain networks that control these functions (Liu, 2016; Iacovella and Hasson, 2011), it will be important to distinguish between spatially widespread effects that may be largely considered as measurement confounds and more localized effects that may be viewed as meaningful reflections of links between DFC measures and physiological activity. Future DFC studies would benefit from the development of analysis methods that more effectively take into account the origins of the nuisance effects.

## 7. Acknowledgements

This work was partially supported by NIH grant R21MH112155 and a UC San Diego Frontiers of Innovation Scholars Program (FISP) Project Fellowship.

## APPENDIX A. Block regression on windowed time series

For the kth window, let *x*_1,*k*_ and *x*_2,*k*_ be a pair of windowed fMRI time series, with *x*̃_1,*k*_ and *x*̃_2,*k*_ corresponding to the time courses after block regression using *n_k_* as the nuisance regressor. Since each window is treated independently in block regression, we can simplify the derivations by dropping the window index *k* for now. We also assume without loss of generality that the time series have zero mean.

For *x*_1_ we can write the time course after regression as:

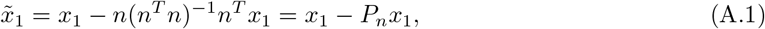

where *P_n_* = *n*(*n^T^n*)^−1^*n^T^* is the projection matrix onto n, and a similar expression holds for *x*_2_. For the derivations that follow, we decompose the nuisance regressor into in-plane and orthogonal components such that *n* = *n_I_* + *n_O_* where *n_I_* is the component of *n* that lies in the subspace spanned by *x*_1_ and *x*_2_, and *n_O_* is the orthogonal complement.

The squared norm for the time series after regression is:

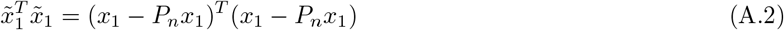

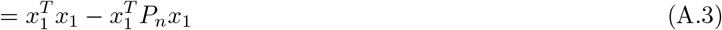

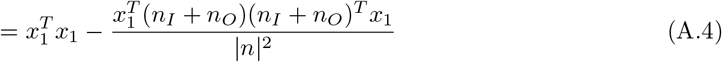

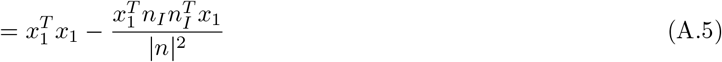

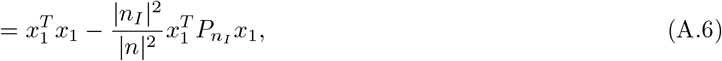

where 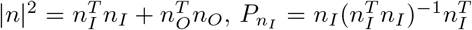, and we have made use of the relation 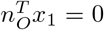 and the symmetry of the projection matrices. The corresponding norm can then be written as:

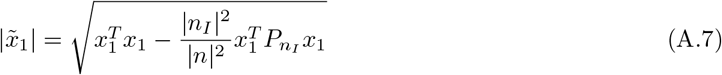

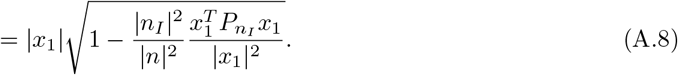

A similar derivation holds for |*x*̃_2_|.

To compute the correlation coefficient after regression, we start with the dot product between *x*̃_1_ and *x*̃_2_ and use the orthogonal decomposition to write:

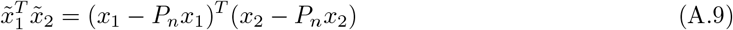

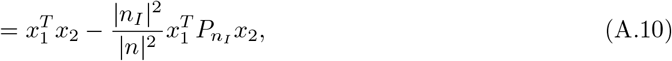

where the omitted steps in the derivation are similar to those shown above for the derivation of the norm. Normalizing the dot product by the appropriate norms yields the correlation coefficient after regression

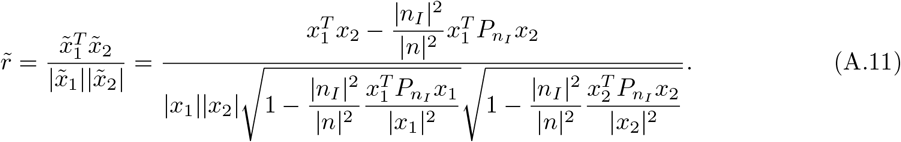

This can be rewritten in the following form:

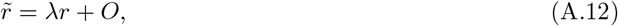

where 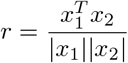 is the correlation coefficient prior to regression, and the scaling λ and offset *O* terms are defined as:

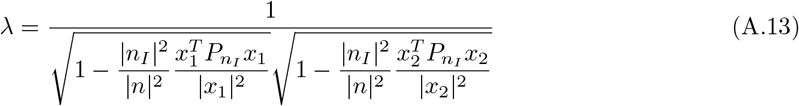

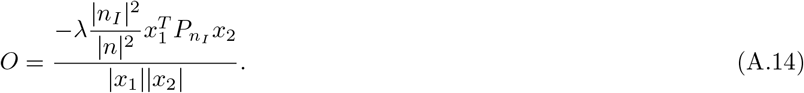

Note that if the orthogonal nuisance component *n*_O_ is large compared to the in-plane component *n_I_*, the ratio 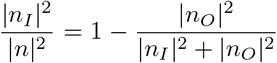 approaches 0, such that the scaling term λ approaches 1, and the offset term *O* goes to 0. As a result, the correlation coefficients before and after regression will be approximately equal *r*̃ ≈ *r*.

To gain further insight, it is useful to rewrite Equation (A.11) using trigonometric functions. First, note that the correlation coefficient prior to regression can be also written as *r* = cos*θ*, where *θ* is the angle between *x*_1_ and *x*_2_. Then without loss of generality, we can write *θ* = *θ*_1_ + *θ*_2_ where *θ*_1_ is the angle between *n_I_* and *x*_1_ and *θ*_2_ is the angle between *n_I_* and *x*_2_, where both *θ*_1_ and *θ*_2_ are assumed to be non-negative. This corresponds to the case where the in-plane term *n_I_* “lies” between the vectors *x*_1_ and *x*_2_. Note it also possible for the in-plane term to lie outside of the vectors, such that *θ* = ±(*θ*_1_ – *θ*_2_). We refer to this second case as the complementary case.

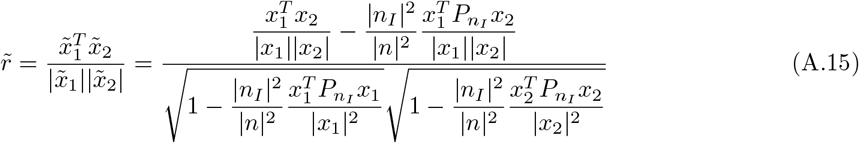

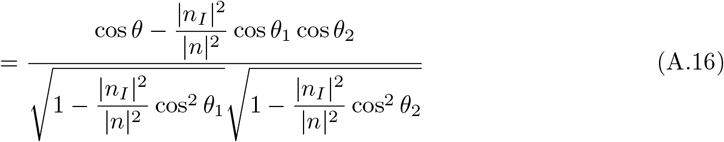

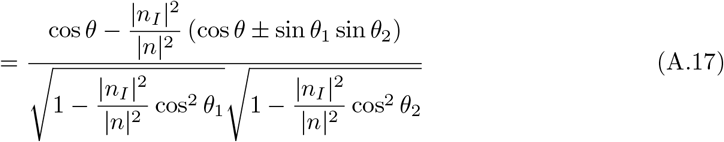

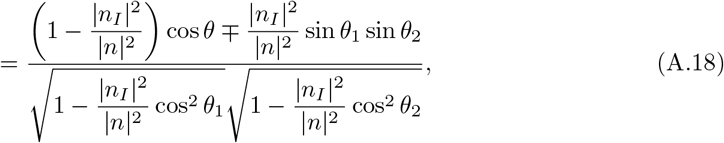

where we have used the identities:

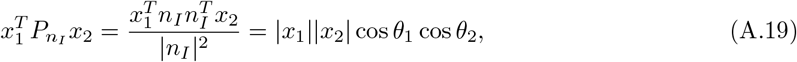

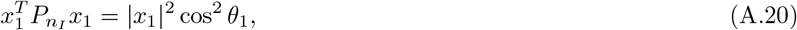

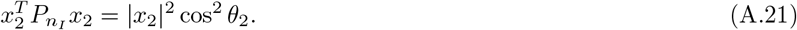

Also note that in Equation (A.17) we have used the trigonometric identity cos*θ*_1_ cos*θ*_2_ = cos*θ* ± sin*θ*_1_ sin*θ*_2_. If *θ* = *θ*_1_ + *θ*_2_, the sign of ∓ in Equation (A.18) is a minus (−), and in the complementary case for *θ* = ±(*θ*_1_ – *θ*_2_) (i.e. when *n_I_* lies outside of *x*_1_ and *x*_2_) the sign is a +.

Finally, by making the substitution 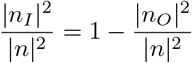, the correlation coefficient after regression is:

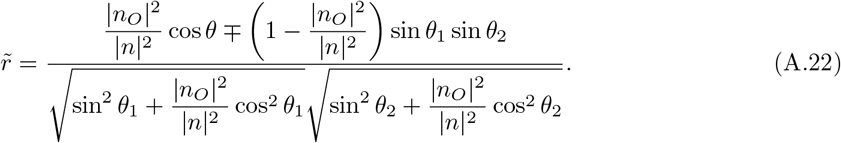

If the orthogonal component becomes small and the orthogonal fraction 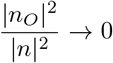, then *r*̃ reduces to:

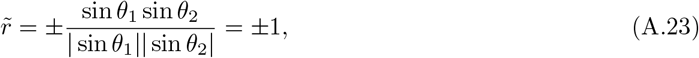

where the minus sign applies to the case where the nuisance vector lies between the observed vectors and the plus sign applies to the complementary case where the nuisance vector lies outside the vectors.

## APPENDIX B. Derivation of limits on difference in DFC estimates

Here we derive the analytical bound on the difference ΔDFC = *r*̃ – *r*, between the pre and post regression DFC estimates. First, we provide derivations for the case when *θ* = *θ*_1_ + *θ*_2_ (we will consider the complementary case *θ* = ±(*θ*_1_ – *θ*_2_) later). In this case, the ΔDFC is:

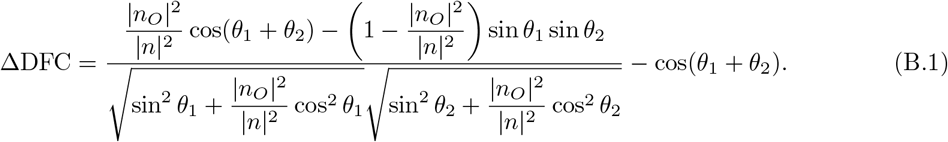

Note that since n¡ lies between the observed vectors *x*_1_ and *x*_2_, nuisance regression will increase the angle (and decrease the correlation value) between the vectors *x*_1_ and *x*_2_, and we will have *r*̃ < *r*. As a result, we are looking for a lower bound on ΔDFC as a function of 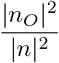.

It can be shown empirically that the expression for ΔDFC is minimized when *θ*_1_ = *θ*_2_. Using this, we simplify the original expression for ΔDFC in Equation B.1 as:

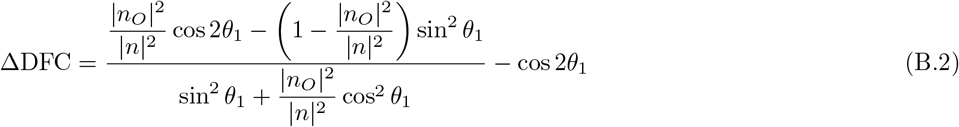

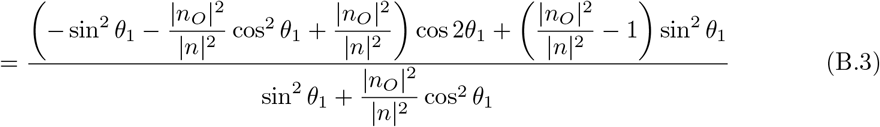

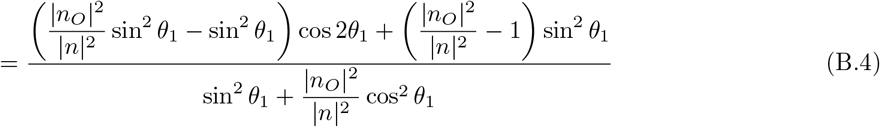

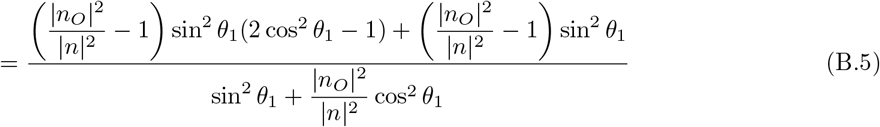

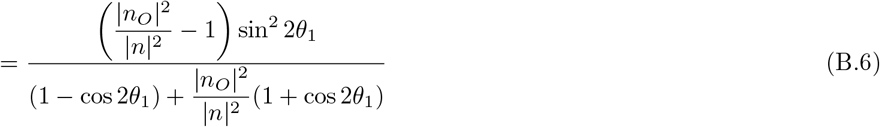

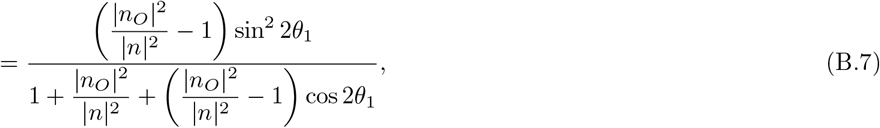

where in the numerator of Equation (B.6), we used the trigonometric identity: 2 sin^2^ *θ*_1_ cos^2^ *θ*_1_ = sin^2^ 2*θ*_1_/2, and in the denominator of the same equation, we used: sin^2^ *θ*_1_ = (1 – cos2*θ*_1_)/2 and cos^2^ *θ*_1_ = (1 + cos2*θ*_1_)/2. We then take the partial derivative with respect to *θ*_1_ as:

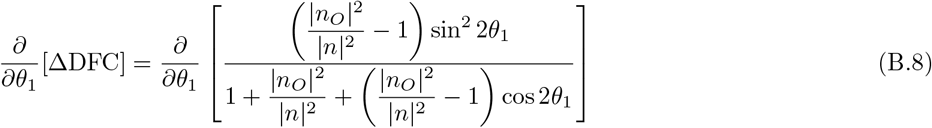

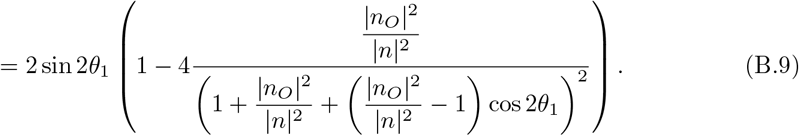

Now solving for the non-trivial roots of the derivative yields 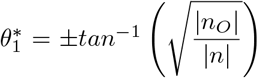. Finally, by using this value in the ΔDFC expression we find the lower bound as:

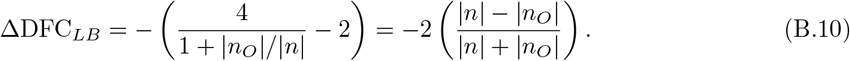

For the complementary case: *θ* = ±(*θ*_1_ – *θ*_2_), in which *n_I_* is positioned outside of *x*_1_ and *x*_2_, linear regression will decrease the angle between the vectors *x*_1_ and *x*_2_ and the correlation value after regression will increase yielding ΔDFC ≥ 0. Therefore, we are seeking an upper bound. It can be shown empirically that the expression for ΔDFC is maximized when *θ*_1_ + *θ*_2_ = π. Following a derivation similar to the one used for the lower bound, the upper bound is obtained as:

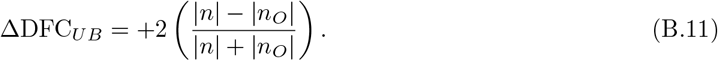

## APPENDIX C. Derivation of DFC estimate after full regression

In this section, we show that the correlation coefficients obtained after full regression can be expressed as a modified version of the correlation coefficients obtained with block regression. First, we note that for the kth window the residual after block regression can be written as:

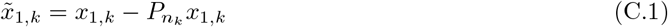

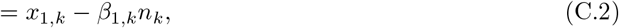

where 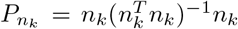 is the projection matrix onto the windowed nuisance time series *n_k_*, and 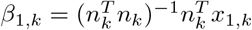 is the corresponding scalar fit coefficient computed for the windowed time series *x*_1,*k*_.

^x^1,k.

For full regression the residual can be written as:

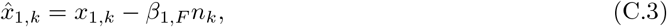

where *β*_1,*F*_ = (*n^T^n*)^−1^ *n^T^ x*_1_ is the scalar fit coefficient computed using the entire time series x_1_ and the full nuisance time series *n*.

Noting that the expressions in Equation (C.2) and Equation (C.3) differ only in the scalar fit coefficients *β*_1,*F*_ and *β*_1,*k*_, we can rewrite Equation (C.3) as:

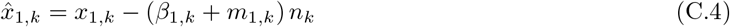

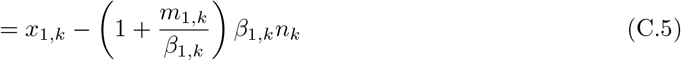

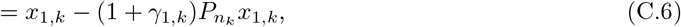

where *m*_1,*k*_ = *β*_1,*F*_ – *β*_1,*k*_ is the difference in fit coefficients and 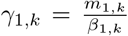 is a correction term that captures the difference between full and block regression. Note that if *γ*_1,*k*_ = 0 then full regression and block regression have identical effects for the kth window. The expression for *x*̂_2,*k*_ has the same form as Equation (C.6).

To compute the correlation coefficient between *x*̂_1,*k*_ and *x*̂_2,*k*_, we first note that the dot product can be written as:

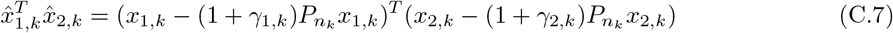

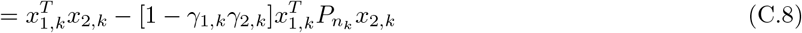

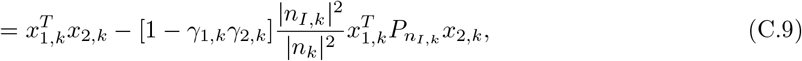

which is similar in form to the dot product expression previously derived in Equation (A.10) for block regression with the addition of a correction term [1 – *γ*_1,*k*_ *γ*_2,*k*_] and the window index *k*. The norms |*x*̂_1,*k*_| and |*x*̂_1,*k*_| can be derived using similar modifications.

Incorporating all the correction terms, we can write the correlation coefficient after full regression as:

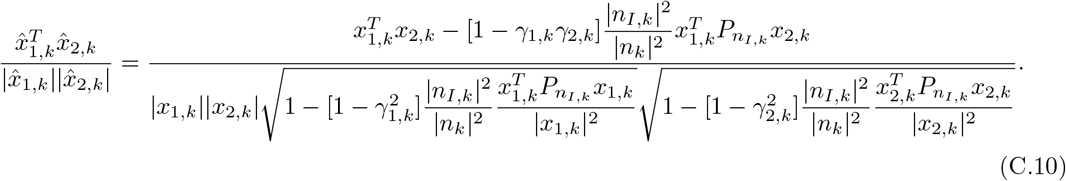

Note that if *γ*_1,*k*_ = *γ*_2,*k*_ = 0 and we drop the window index k, then this expression is identical to the expression for block regression previously shown in Equation (A.11).

## APPENDIX D. Nuisance Norm Regression (NNR)

Nuisance norm regression aims to remove the nuisance norm-related variance from the DFC estimates. In this technique, nuisance removal can be performed in the relevant DFC metric space in addition to traditional nuisance removal on raw time series. For sliding window correlations obtained between pairs of ROIs, we define a sliding window correlation vector *R* = [*r*_1_, *r*_2_, ∆, *r_k_*, ∆, *r_M_*]^*T*^, where *k* is the window index, *M* is the total number of windows, and *r_k_* is the pairwise correlation value for the *k*th window. Similarly, we define a nuisance norm vector (or matrix for multiple nuisance terms) *N* = [|*n*_1_|, |*n*_2_|, ∆, |*n_k_*|, ∆, |*n_M_*|]^*T*^, where |*n_k_*| is the norm of the nuisance regressor for the kth window. Then the effects of a nuisance norm can be removed through linear regression to obtain a “clean” DFC estimate *R* = *R* – *N*(*N^T^ N*)^−^ *N^T^ R*. In the case of multiple nuisance terms, the nuisance norm matrix can be expanded to include additional norm vectors. For example, let *N_WM_*, *N_CSF_*, *N_RV f_* , *N_HR f_*, and *N_HM_* be the sliding window norm vectors for the WM, CSF, RVf, HRf, and HM (total norm) regressors, respectively. Then the corresponding nuisance norm matrix is *N* = [*N_WM_*, *N_CSF_*, *N_RV f_*, *N_HRf_*, *N_HM_*]. Potential variations include expansion of the nuisance norm matrix to include additional terms, such as the squared norms or temporal derivatives of the sliding window norms.

---

**Supplementary Table 1:** Percentage of scans exhibiting significant correlations between the DFC estimates and nuisance norms across all the cases examined. We show the results for window sizes of ~40 seconds, ~58 seconds and ~100 seconds as indicated in the top row. The significance thresholds are indicated in the leftmost column. For each significance level, the percentage of scans with significant correlations are shown for the Pre (prior to nuisance regression), Post (after full regression), and BlockReg (after block regression) conditions. The columns with the ‘Overall’ label show the significance results when considering all regressors. The columns with the ‘GS’ label display the percentage of scans with significant correlations when considering only the GS regressor. The columns with the label ‘Non-GS’ show the significance results when considering all regressors except for the GS.

| Significance Level | State | % of Significant Scans<br>(Window Size: 40 seconds) |  |  | % of Significant Scans<br>(Window Size: 58 seconds) |  |  | % of Significant Scans<br>(Window Size: 100 seconds) |  |  |
| --- | --- | --- | --- | --- | --- | --- | --- | --- | --- | --- |
|  |  | *GS* | *Non-GS* | ***Overall*** | *GS* | *Non-GS* | ***Overall*** | *GS* | *Non-GS* | ***Overall*** |
| $p \leq 0.05$ | Pre | 52% | 15% | **24%** | 44% | 17% | **24%** | 43% | 24% | **29%** |
|  | Post | 12% | 11% | **11%** | 12% | 15% | **14%** | 20% | 24% | **23%** |
|  | BlockReg | 11% | 11% | **11%** | 11% | 15% | **14%** | 18% | 24% | **22%** |
| $p \leq 0.10$ | Pre | 59% | 23% | **32%** | 52% | 23% | **30%** | 53% | 37% | **41%** |
|  | Post | 21% | 20% | **20%** | 19% | 22% | **22%** | 29% | 35% | **33%** |
|  | BlockReg | 20% | 19% | **20%** | 19% | 23% | **22%** | 29% | 33% | **32%** |

**Supplementary Figure 1:**
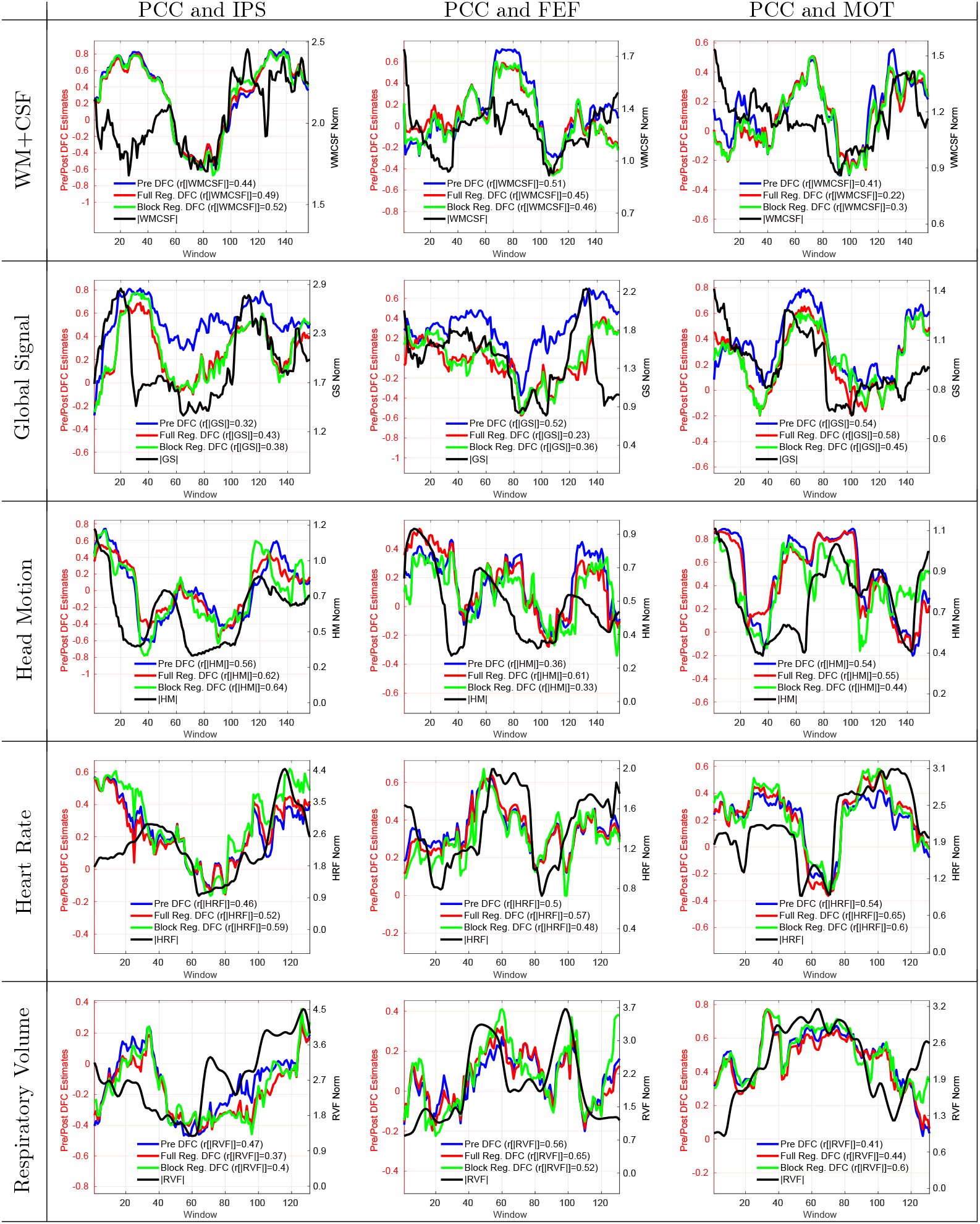
15 example scans that demonstrate a range of moderate to strong correlations (0.30 < *r* < 0.65) between the nuisance norms and the DFC estimates. The type of nuisance regressor is indicated by the row label. The seed pair for the DFC estimate is indicated by the column label. There is some evidence of a delay between the nuisance norm and DFC estimates as in the first two columns for the HM regressors.

**Supplementary Figure 2:**
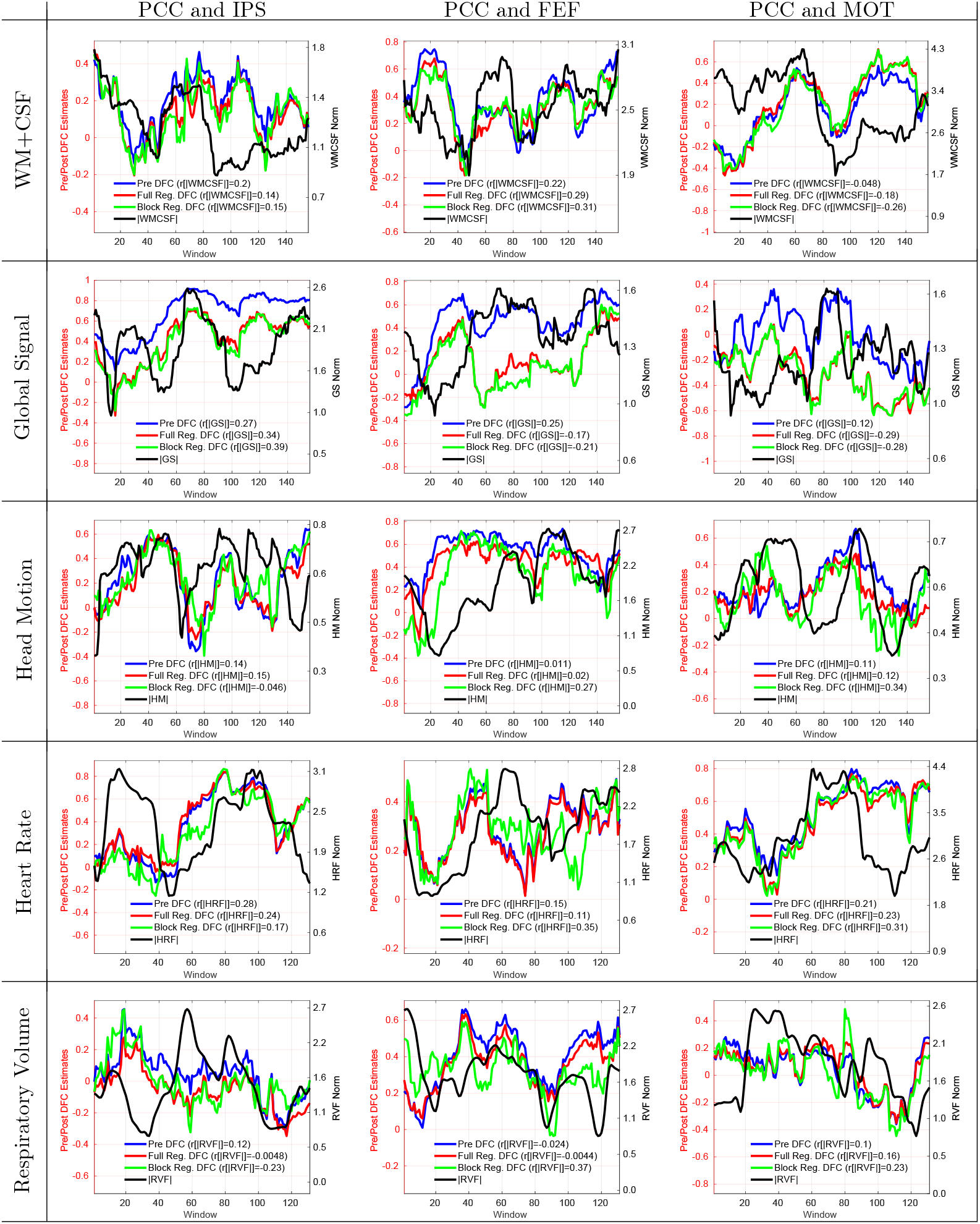
15 example scans that demonstrate weaker correlation values (*r* < 0.30) between the nuisance norms and the Pre DFC estimates. The type of nuisance regressor is indicated by the row label. The seed pair for the DFC estimate is indicated by the column label. Even though the observed correlations are weaker in this Figure, there is some visual similarity between the nuisance norms and DFC estimates.

**Supplementary Figure 3:**
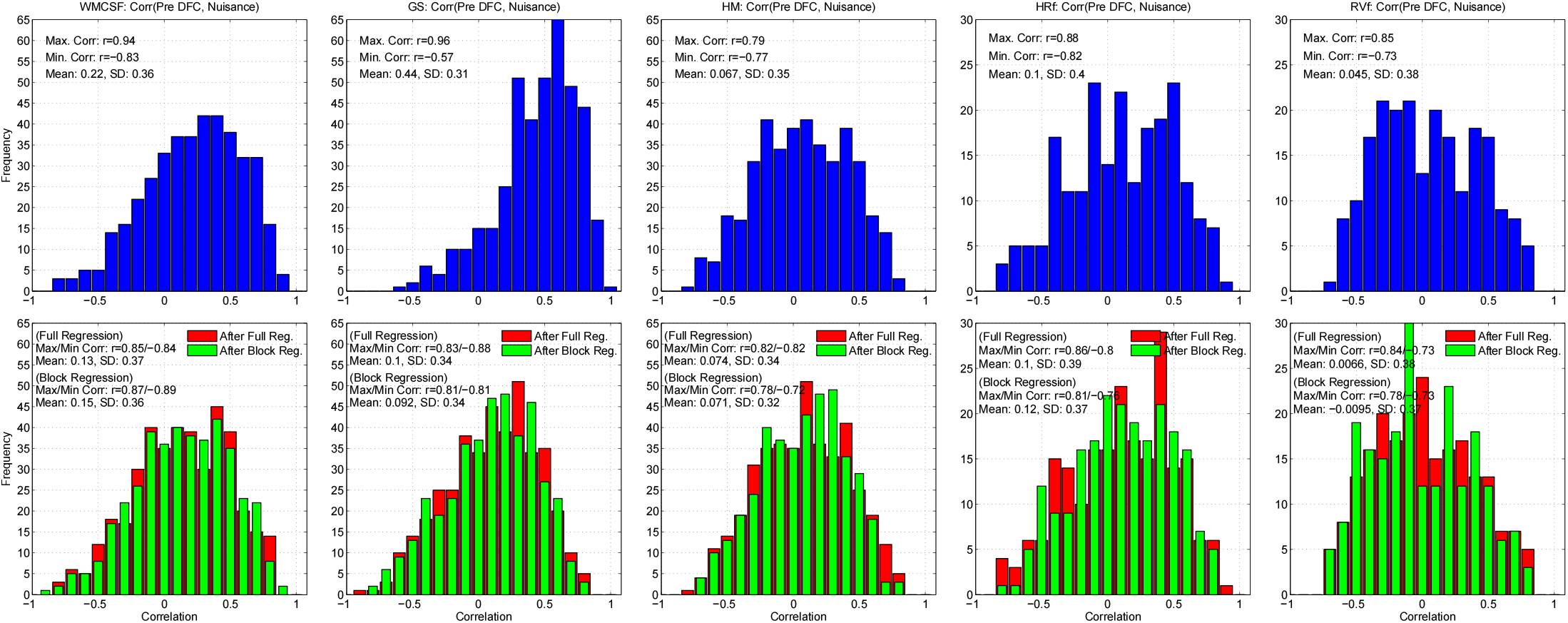
This figure shows the correlations presented in Figure 3 grouped according to individual regressor norms. The first row in each column shows the histogram of correlation values obtained between the Pre DFC estimates and the nuisance norm across all scans, seed pairs for a specific nuisance regressor (columns from left to right correspond to WM+CSF, GS, HM, HRF, and RVf norms). The second row shows the correlation values after full/block regression for different regressors. The minimum and maximum correlations as well as the mean correlation value for each group is labeled in each subplot.

**Supplementary Figure 4:**
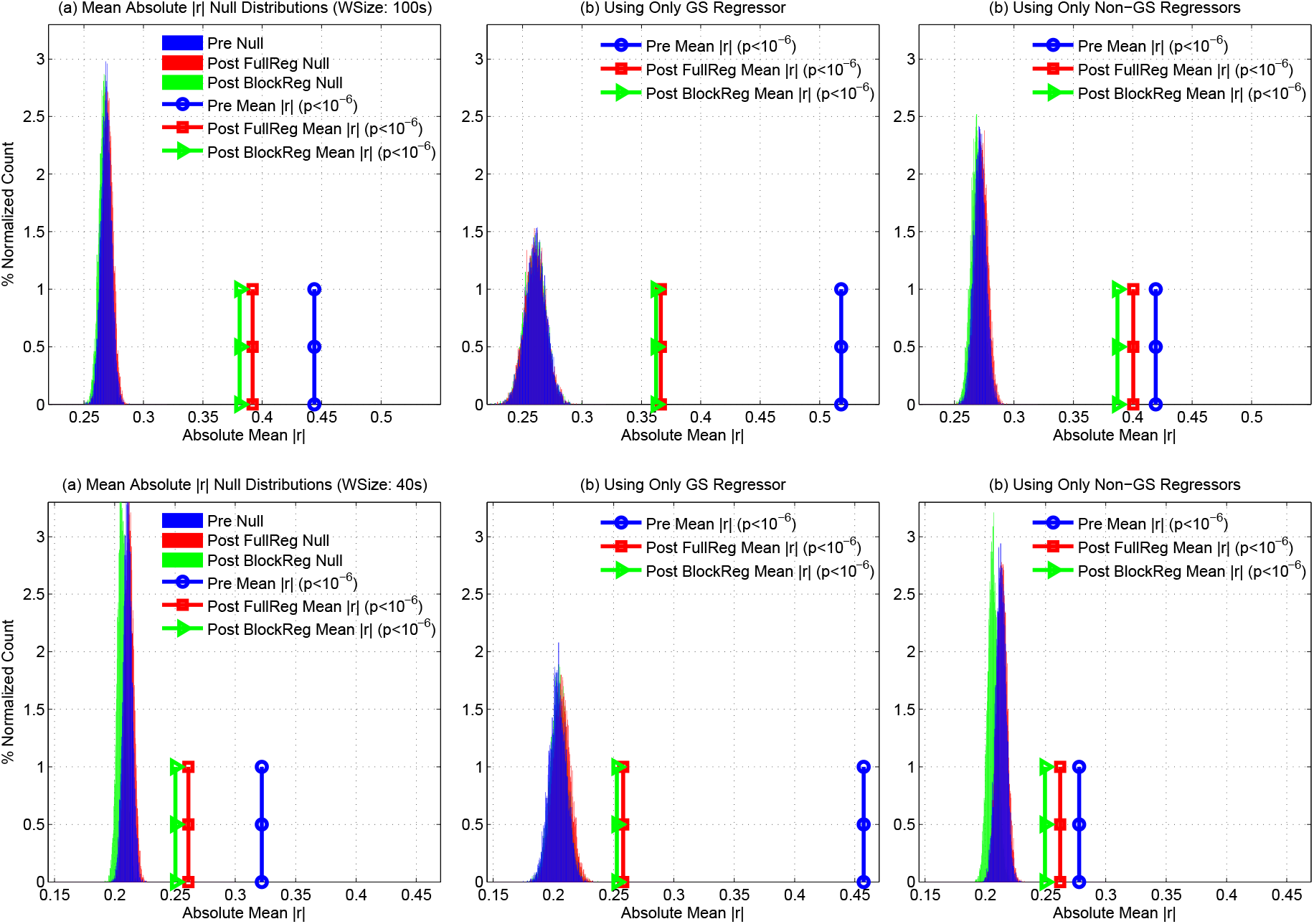
Empirical null distributions and sample mean absolute correlation values for window sizes of 100s (top row) and 40s (bottom row). (a) Empirical null distributions for mean absolute correlation values when considering all regressors both before regression (blue) and after full regression (red) or block regression (green). The sample mean absolute correlation values for these conditions are indicated by the circles, squares, and triangles. These values were found to be significant *p* < 10^−6^ both before and after full or block regression. (b) Empirical null distributions and sample mean absolute correlation values when considering only the GS regressor. There was a marked reduction in the sample mean absolute correlation values after regression. However, the sample mean absolute correlation values were still found to be significant *p* < 10^−6^ both before and after nuisance regression. (c) Empirical null distributions and sample mean absolute correlation values for the Non-GS regressors. Sample mean absolute correlation values were found to be significant *p* < 10^−6^ both before and after nuisance regression

**Supplementary Figure 5:**
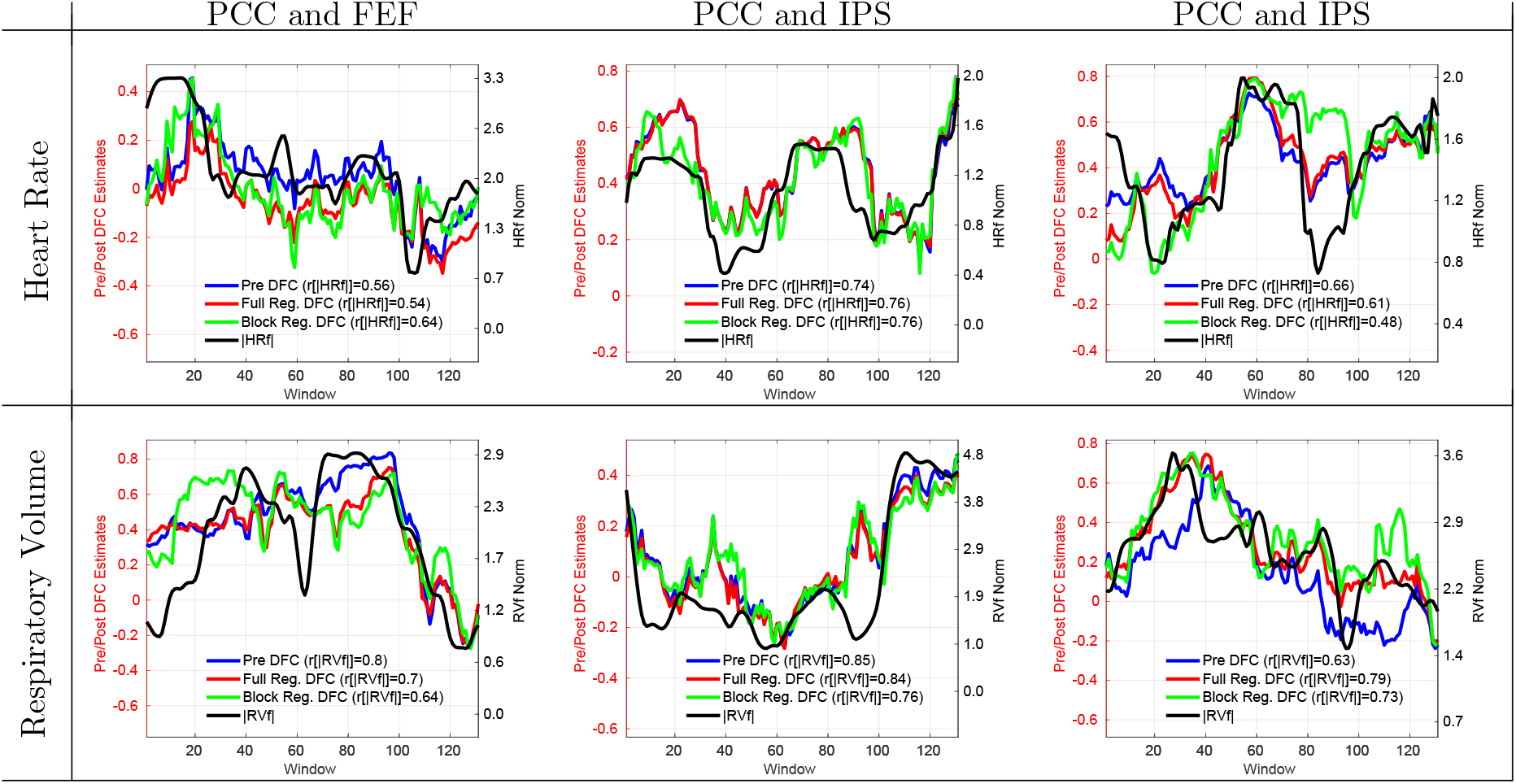
6 representative scans showing that both before and after nuisance regression the DFC estimates (e.g. between PCC and IPS, PCC and MOT and PCC and FEF) are strongly correlated with the respective nuisance norms.

**Supplementary Figure 6:**
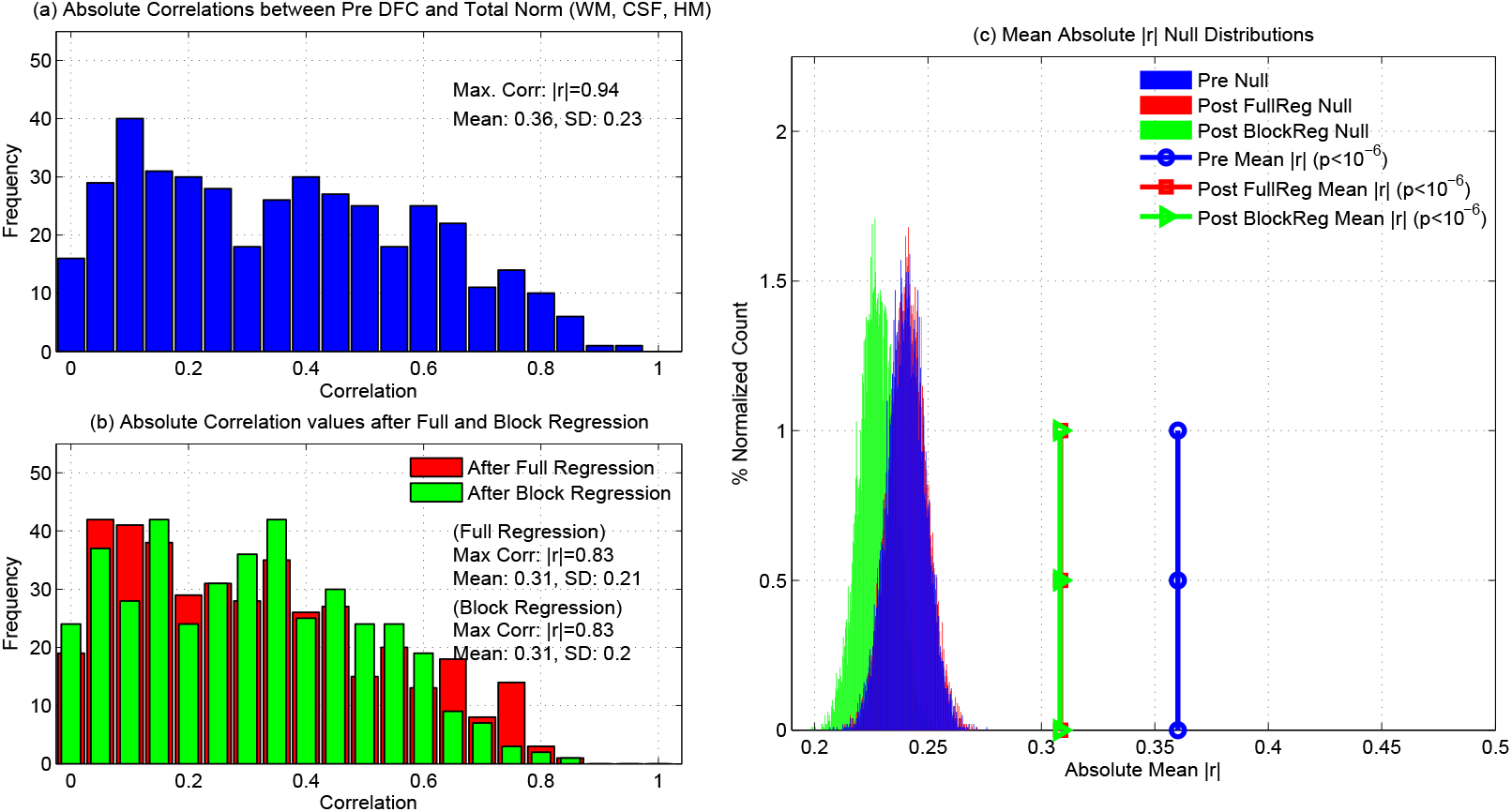
Results obtained when using WM, CSF, and 6 HM regressors for nuisance regression. (a) Histogram of absolute correlations (before nuisance regression) between the total norm of the WM, CSF and 6 HM regressors and Pre DFC estimates obtained across all seed pairs. The mean of the absolute correlations across all cases was |*r*| = 0.36. The signed correlation values (not shown) ranged from *r* = −0.83 to *r* = 0.94 with a skewed distribution in which 72% of the correlations were positive and the remaining 28% were negative. (b) Histogram of absolute correlations between the total norm after full and block regression with sample means for both cases of |*r*| = 0.31. The signed correlations (not shown) ranged from *r* = −0.80 to *r* = 0.83 for full regression (65% of the correlations were positive) and from *r* = −0.75 to *r* = 0.83 for block regression (69% of the correlations were positive). The distribution of absolute correlation values before and after nuisance regression were similar with cosine similarity values of *S* = 0.82 and *S* = 0.75 for full and block regression, respectively. (c) Empirical null distributions and sample mean absolute correlation values before and after nuisance regression. All sample mean absolute correlation values were significant (*p* < 10^−6^) as assessed using their respective null distribtuions.

**Supplementary Figure 7:**
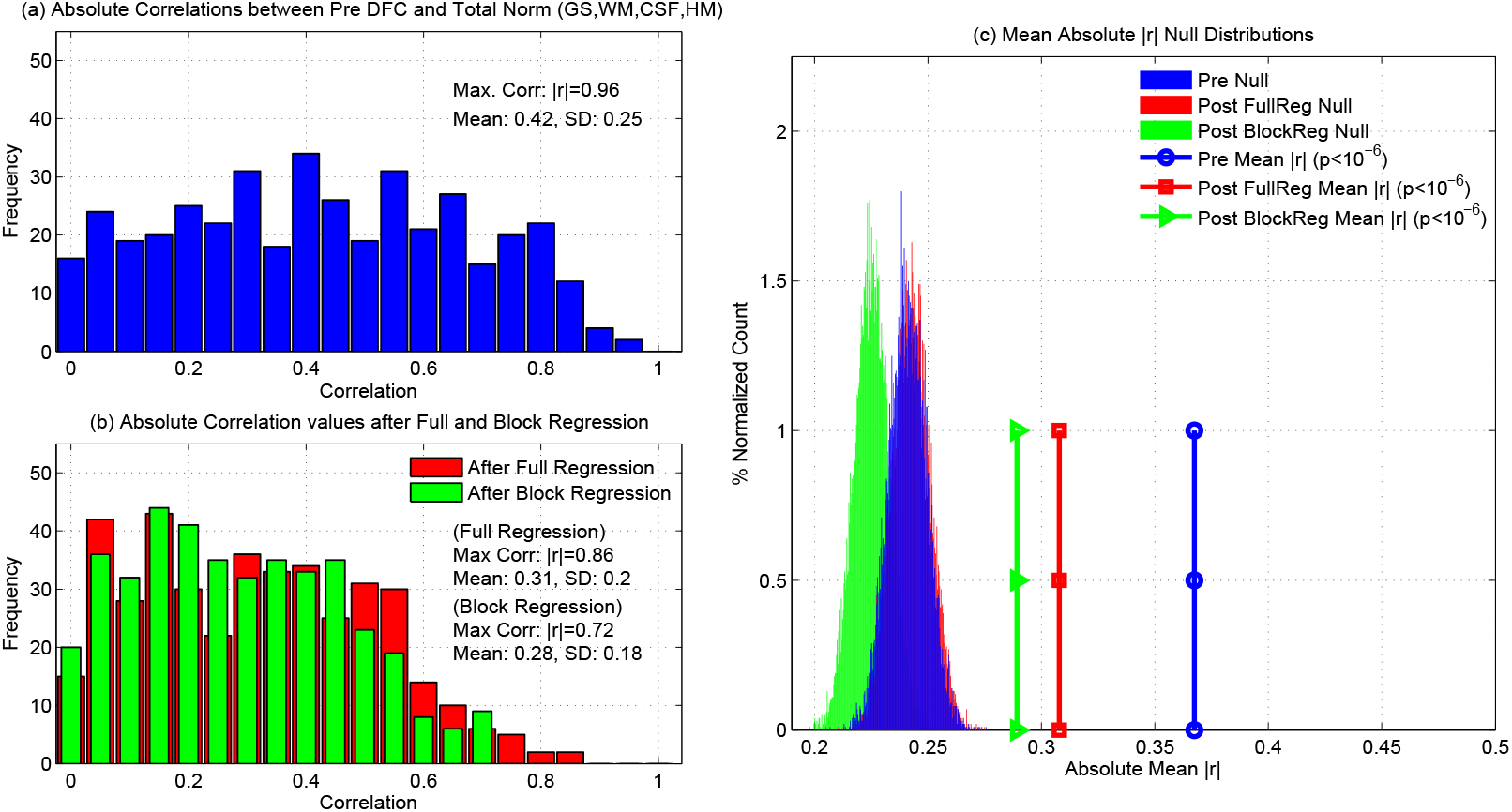
Results obtained when using GS, WM, CSF, and 6 HM regressors for nuisance regression. (a) Histogram of absolute correlations (before nuisance regression) between the total norm of GS, WM, CSF and the 6 HM regressors and Pre DFC estimates obtained across all seed pairs. The mean of the absolute correlations across all cases was |*r*| = 0.42. The signed correlation values (not shown) ranged from *r* = −0.74 to *r* = 0.95 with a skewed distribution in which 96% of the correlations were positive and the remaining 4% were negative. (b) Histogram of absolute correlations after nuisance regression with sample means of |*r*| = 0.31 and |*r*| = 0.29 for full and block regression, respectively. The signed correlations (not shown) ranged from *r* = −0.77 to *r* = 0.86 for full regression (63% of the correlations were positive) and from *r* = −0.72 to *r* = 0.71 for block regression (67% of the correlations were positive). The distribution absolute correlation values before and after multiple regression were similar with a cosine similarity value of *S* = 0.79 for both full and block regression. (c) Empirical null distributions and sample mean absolute correlation values before and after nuisance regression. All sample mean absolute correlation values were significant (*p* < 10^−6^) as assessed using their respective null distributions.

**Supplementary Figure 8:**
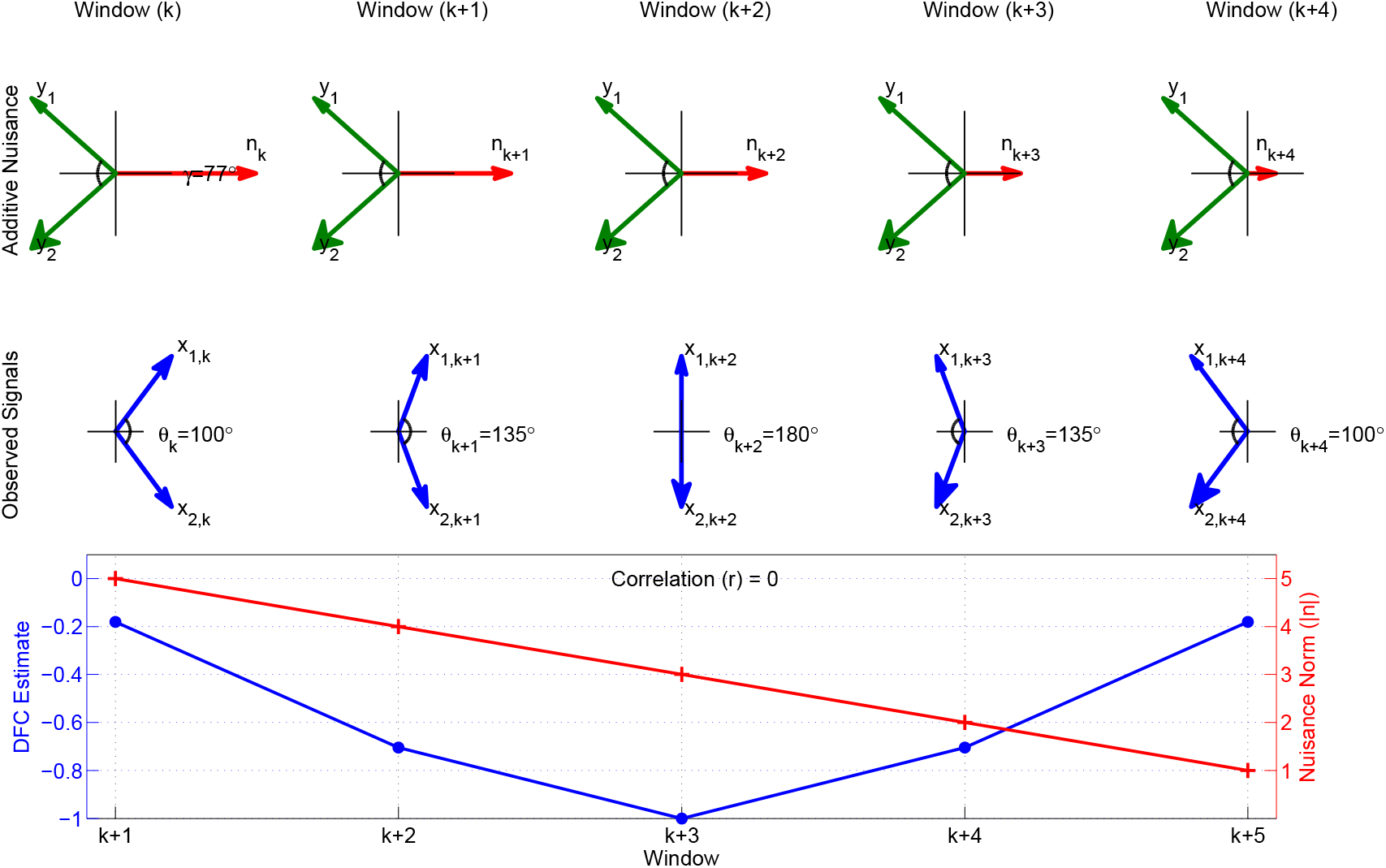
In the first three windows (*k, k* + 1, and *k* + 2) a decrease in the nuisance norm corresponds to a decrease in the DFC estimates since the angle between the observed vectors increases as the norm decreases. In the last two windows (*k* + 3 and *k* + 4), the nuisance norm continues to decrease and a crossover occurs such that the angle between the observed vectors starts to decrease, leading to an increase in the DFC estimates. Overall, the nuisance term induces a decrease in DFC estimates in the first 3 windows and an increase in the last two windows. Thus the nuisance norm appears uncorrelated with the DFC estimates with *r* = 0, even though the nuisance term has a clear effect on the observed vectors.

**Supplementary Figure 9:**
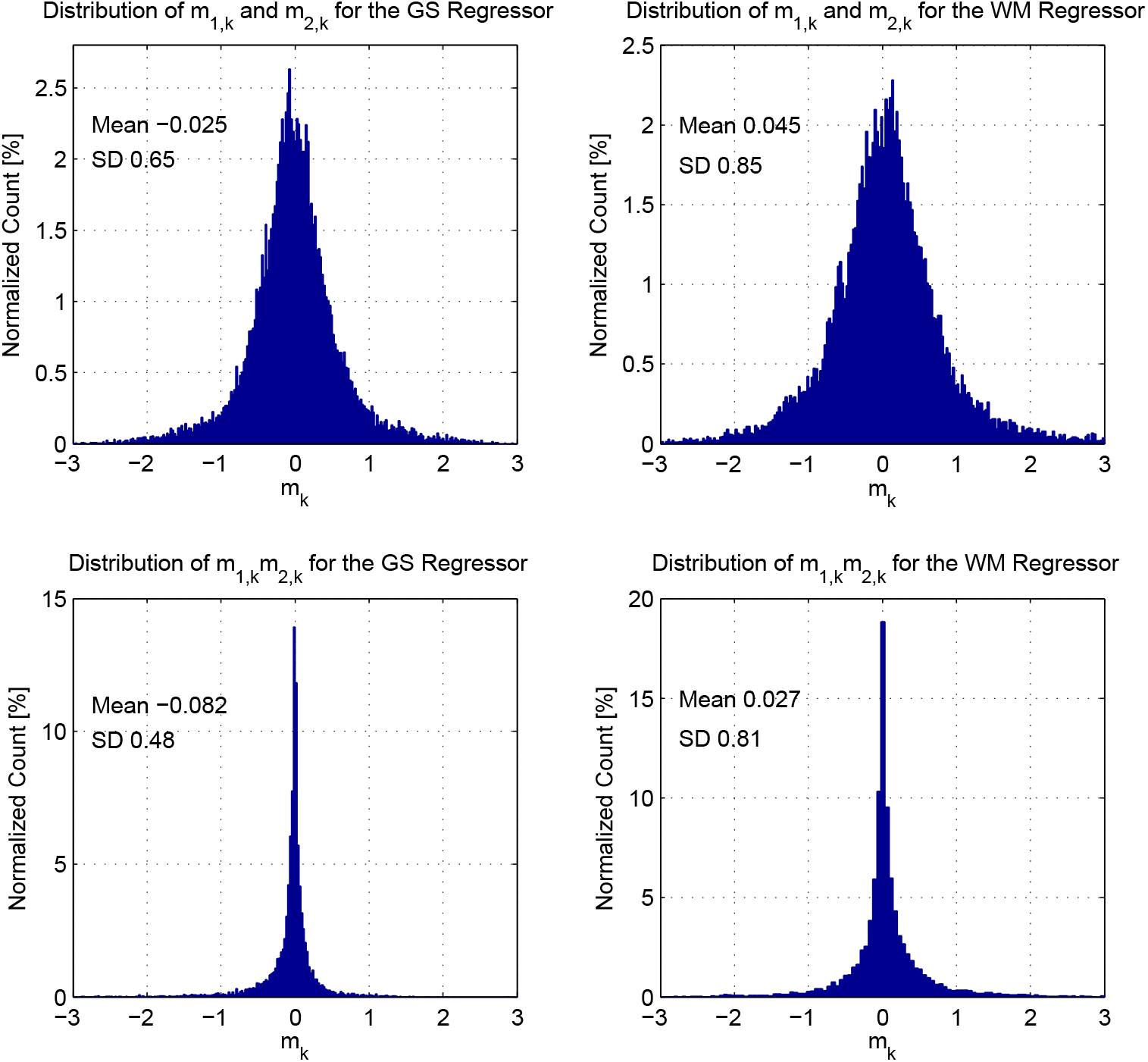
Top row: Histograms of the difference terms *m*_1,*k*_ = *β*_1,*F*_ − *β*_1,*k*_ and *m*_2,*k*_ = *β*_2,*F*_ − *β*_2,*k*_ for GS and WM regressors. The plots combine the instances of the two terms. Bottom row: Histograms of the product term *m*_1,*k*_*m*^2,*k*^.

**Supplementary Figure 10:**
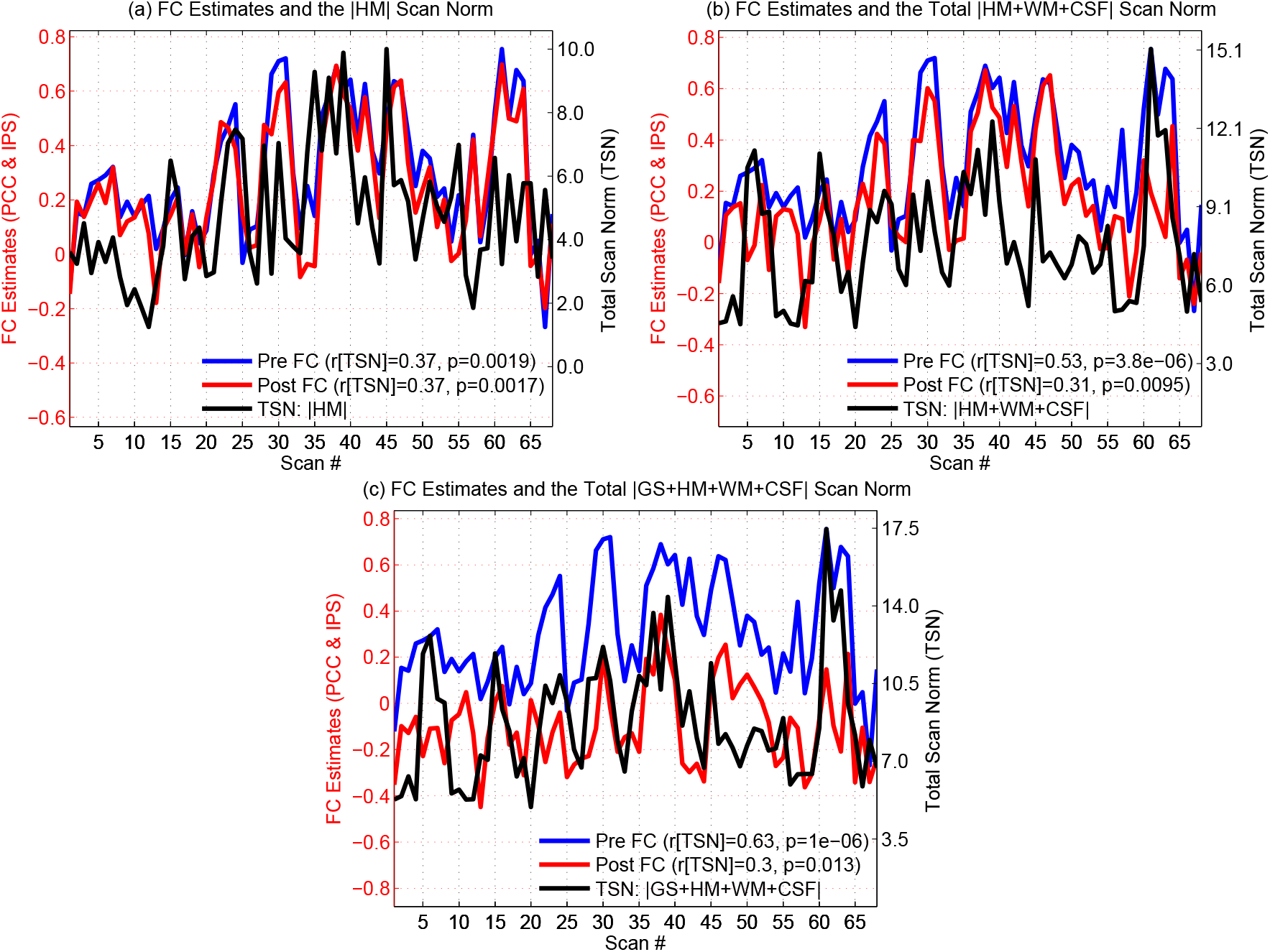
Example of relation between static FC estimates and nuisance norms. In each of the panels, the blue line shows the per-scan estimates (prior to nuisance regression) of static FC between the PCC and IPS seeds versus the scan number. The black lines in each of the panels shows the per-scan nuisance norm as a function of scan number, where the nuisance terms used in computing the total nuisance norms were (a) 6 HM regressors, (b) WM, CSF, and 6 HM regressors, and (c) GS, WM, CSF, and 6 HM regressors. For each combination of regressors, the static FC estimates were significantly correlated (*r* > 0.37;*p* < 2 × 10^−3^) across scans with the total nuisance norms, with the correlation coefficients and p-values for each case indicated in each panel. The red lines in each of the panels show the static FC estimates after performing nuisance regression using the regressors previously described for each panel. After regression, the static FC estimates were still significantly correlated (*r* > 0.30; *p* < 0.02) across scans with the total nuisance norms, with the correlation coefficients and p-values for each case indicated in each panel.

